# Price equation captures the role of drug interactions and collateral effects in the evolution of multidrug resistance

**DOI:** 10.1101/2020.11.06.371682

**Authors:** Erida Gjini, Kevin B. Wood

## Abstract

Bacterial adaptation to antibiotic combinations depends on the joint inhibitory effects of the two drugs (drug interaction, DI) and how resistance to one drug impacts resistance to the other (collateral effects, CE). Here we model these evolutionary dynamics on two-dimensional phenotype spaces that leverage scaling relations between the drug-response surfaces of drug sensitive (ancestral) and drug resistant (mutant) populations. We show that evolved resistance to the component drugs–and in turn, the adaptation of growth rate–is governed by a Price equation whose covariance terms encode geometric features of both the two-drug response surface (DI) in ancestral cells and the correlations between resistance levels to those drugs (CE). Within this framework, mean evolutionary trajectories reduce to a type of weighted gradient dynamics, with the drug interaction dictating the shape of the underlying landscape and the collateral effects constraining the motion on those landscapes. Our results clarify the complex relationship between drug interactions and collateral effects in multi-drug environments and illustrate how specific dosage combinations can shift the weighting of these two effects, leading to different and temporally-explicit selective outcomes.

## Introduction

Understanding and predicting evolutionary dynamics is an ongoing challenge across all fields of biology. Microbial populations offer relatively simple model systems for investigating adaptation on multiple length scales, from the molecular to the population level, on timescales ranging from a few generations to decades. Yet even these simplest of systems exhibit rich and often counter intuitive evolutionary dynamics. Despite enormous progress, both theoretical and experimental, predicting evolution remains exceedingly difficult, in part because it is challenging to identify the phenotypes, selective gradients, and environmental factors shaping adaptation. In turn, controlling those dynamics-for example, by judicious manipulation of environmental conditions-is often impossible. These challenges represent open theoretical questions but also underlie practical public health threats–exemplified by the rapid rise of antibiotic resistance (***Davies and Davies, 2010***; ***Levy and Marshall, 2004***)–where evolutionary dynamics are fundamental to the challenge, and perhaps, to the solution.

In recent years, evolution-based strategies for impeding drug resistance have gained significant attention. These approaches have identified a number of different factors that could modulate resistance evolution, including spatial heterogeneity (***Zhang et al., 2011***; ***Baym et al., 2016a***; ***Greulich et al., 2012***; ***Hermsen et al., 2012***; ***Moreno-Gamez et al., 2015***; ***Gokhale et al., 2018***; ***De Jong and Wood, 2018***; ***Santos-Lopez et al., 2019***); competitive (***Read et al., 2011***; ***Hansen et al., 2017, 2020***), cooperative (***Meredith et al., 2015b***; ***Artemova et al., 2015***; ***Sorg et al., 2016***; ***Tan et al., 2012***; ***Karslake et al., 2016***; ***Yurtsev et al., 2016***; ***Hallinen et al., 2020***), or metabolic (***Adamowicz et al., 2018, 2020***) interactions between bacterial cells; synergy with the immune system (***Gjini and Brito, 2016***); epistasis between resistance mutations (***Trindade et al., 2009***; ***Borrell et al., 2013***; ***Lukačišinová et al.,2020***); plasmid dynamics (***Lopatkin et al., 2016, 2017***; ***Cooper et al., 2017***); precise tuning of drug doses (***Lipsitch and Levin, 1997***; ***Yoshida et al., 2017***; ***Meredith et al., 2015a***; ***Nichol et al., 2015***; ***Fuentes-Hernandez et al., 2015***; ***Coates et al., 2018***; ***Iram et al., 2020***); cycling or mixing drugs at the hospital level (***Bergstrom et al., 2004***; ***Beardmore et al., 2017***); and statistical correlations between resistance profiles for different drugs (***Imamovic and Sommer, 2013***; ***Kim et al., 2014***; ***Pál et al., 2015***; ***Barbosa et al., 2017, 2018***; ***Rodriguez de Evgrafov et al., 2015***; ***Nichol et al., 2019***; ***Podnecky et al., 2018***; ***Imamovic et al., 2018***; ***Barbosa et al., 2019***; ***Rosenkilde et al., 2019***; ***Apjok et al., 2019***; ***Maltas and Wood, 2019***; ***Maltas et al., 2020***; ***Hernando-Amado et al., 2020***; ***Roemhild et al.,2020***).

Drug combinations are a particularly promising approach for slowing resistance (***Baym et al., 2016b***), but the evolutionary impacts of combination therapy remain difficult to predict, especially in a clinical setting (***Podolsky, 2015***; ***Woods and Read, 2015***). Antibiotics are said to interact when the combined effect of the drugs is greater than (synergy) or less than (antagonism) expected based on the effects of the drugs alone (***Greco et al., 1995***). These interactions may be leveraged to improve treatments-for example, by offering enhanced antimicrobial effects at reduced concentrations. But these interactions can also accelerate, reduce, or even reverse the evolution of resistance (***Chait et al., 2007***; ***Michel et al., 2008***; ***Hegreness et al., 2008***; ***Pena-Miller et al., 2013***; ***Dean et al., 2020***), leading to trade-offs between short-term inhibitory effects and long-term evolutionary potential (***Torella et al., 2010***). In addition, resistance to one drug may be associated with modulated resistance to other drugs. This cross-resistance (or collateral sensitivity) between drugs in a combination has also been shown to significantly modulate resistance evolution (***Barbosa et al., 2018***; ***Rodriguez de Evgrafov et al., 2015***; ***Munck et al., 2014***).

Predicting these evolutionary dynamics is difficult. Despite notable recent progress, collateral effects (***Pál et al., 2015***; ***Roemhild et al., 2020***) and drug interactions (***Bollenbach et al., 2009***; ***Chevereau et al., 2015***; ***Lukačišin and Bollenbach, 2019***; ***Chevereau et al., 2015***), even in isolation, reflect interactions-between genetic loci, between competing evolutionary trajectories, between chemical stressors-that are often poorly understood at a mechanistic or molecular level. Yet adaptation to a drug combination may often reflect both phenomena, with the pleiotropic effects that couple resistance to individual drugs constraining, or constrained by, the interactions that occur when those drugs are used simultaneously. In addition, the underlying genotype space is high-dimensional and potentially rugged, rendering the genotpyic trajectories prohibitively complex (***De Visser and Krug, 2014***).

In this work, we attempt to navigate these obstacles by modeling evolutionary dynamics on lower-dimensional phenotype spaces that leverage scaling relations between the drug-response surfaces of ancestral and mutant populations. Our approach is inspired by the fact that multi-objective evolutionary optimization may occur on surprisingly low-dimensional phenotypic spaces (***Shoval et al., 2012***; ***Hart et al., 2015***). To develop a similarly coarse-grained picture of multi-drug resistance, we associate selectable resistance traits with changes in effective drug concentrations, formalizing the geometric rescaling assumptions originally pioneered in (***Chait et al., 2007***; ***Hegreness et al.,2008***; ***Michel et al., 2008***) and connecting evolutionary dynamics with a simple measurable property of ancestral populations. We show that evolved resistance to the component drugs-and in turn, the adaptation of growth rate-is governed by a Price equation whose covariance terms encode geometric features of both 1) the two-drug response surface in ancestral populations (the drug interaction) and 2) the correlations between resistance levels to those drugs (collateral effects). In addition, we show how evolutionary trajectories within this framework reduce to a type of weighted gradient dynamics on the two-drug landscape, with the drug interaction dictating the shape of the underlying landscape and the collateral effects constraining the motion on those landscapes, leading to deviations from a simple gradient descent. Our results clarify the complex relationship between drug interactions and collateral effects in multi-drug environments and illustrate how specific dosage combinations can shift the weighting of these two effects, leading to different selective outcomes even when the available genetic routes to resistance are unchanged.

## Results

Our goal is to understand evolutionary dynamics of a cellular population in the presence of two drugs, Drug 1 and Drug 2. These dynamics reflect a potentially complex interplay between drug interactions and collateral evolutionary trade-offs, and our aim is to formalize these effects in a simple model. To do so, we assume that the per capita growth rate of the ancestral population is given by *G*(*x, y*), where *x* and *y* are the concentrations of drugs 1 and 2 respectively. We limit our analysis to two-drug combinations, but it could be extended to higher-order drug combinations, though this would require empirical or theoretical estimates (***Wood et al., 2012***; ***Zimmer et al., 2016, 2017***) for higher-dimensional drug response surface. At this stage, we do not specify the functional form of *g*(*x, y*), though we assume that this function can be derived from pharmacodynamic or mechanistic considerations (***Engelstädter, 2014***; ***Bollenbach et al., 2009***; ***Wood and Cluzel, 2012***) or otherwise estimated from experimental data (***Greco et al., 1995***; ***Wood et al., 2014***). In classical pharmacology models (***Loewe, 1953***; ***Greco et al., 1995***), the shape of these surfaces-specifically, the convexity of the corresponding contours of constant growth (“isoboles”)–determines the type of drug interaction, with linear isoboles corresponding to additive drug pairs. In this framework, deviations from linearity indicate synergy (concave up) or antagonism (concave down). While there are multiple conventions for assigning geometric features of the response surface to an interactions type-and there has been considerable debate about the appropriate null model for additive interactions (***Greco et al., 1995***)-the response surfaces contain complete information about the phenotypic response. The manner in which this response data is mapped to a qualitative interaction type-synergy or antagonism, for example-is somewhat subjective, though in what follows we adopt this classical Loewe approach for clarity.

### Resistance and rescaling in a simple model

The primary assumption of the model is that the phenotypic response (e.g. growth rate) of drug resistant mutants, which may be present in the initial population or arise through mutation, corresponds to a simple rescaling of the growth rate function *G*(*x, y*) for the ancestral population. As we will see, this scheme provides an explicit link between a cell’s level of antibiotic resistance and its fitness in a given multidrug environment. Specifically, we assume that the growth rate (*g_i_*) of mutant *i* is given by

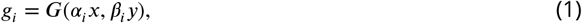

where *α_i_* and *β_i_* are rescaling parameters that reflect the physiological effects of mutations on the growth rate. In some cases-for example, resistance due to efflux pumps (***Wood and Cluzel, 2012***) or drug degrading enzymes (***Yurtsev et al., 2013***)-this effective concentration change corresponds to a physical change in intracellular drug concentration. More generally, though, this hypothesis assumes that resistant cells growing in external drug concentration *x* behave similarly to wild type (drug-sensitive) cells experiencing a reduced effective concentration *αx*. Similar rescaling arguments were originally proposed in (***Chait et al., 2007***), where they were used to predict correlations between the rate of resistance evolution and the type of drug interaction. These arguments have also been used to describe fitness tradeoffs during adaptation (***Das et al., 2020***) and to account for more general changes in the dose response curves, though in many cases the original two-parameter rescaling was sufficient to describe the growth surface in mutants (***Wood et al., 2014***).

When only a single drug is used, this rescaling leads to a simple relationship between the characteristic inhibitory concentrations-for example, the half-maximal inhibitory concentration (IC_50_) or the minimum inhibitory concentration (MIC)-of the ancestral (sensitive) and mutant (resistant) populations. In what follows, we refer to these reference concentrations as 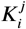, where *i* labels the cell type and, when there is more than one drug, *j* labels the drug. Conceptually, this means that dose response curves for both populations have the same basic shape, with resistance (or sensitivity) in the mutant corresponding only to a shape-preserving rescaling of the drug concentration (*D* → *D/K_i_*; Figure 1A). In the presence of two drugs, the dose response curves become dose response surfaces, and rescaling now corresponds to a shape-preserving rescaling of the contours of constant growth. There are now two scaling parameters, one for each drug, and in general they are not equal. For example, in Figure 1B, the mutant shows increased sensitivity to drug 1 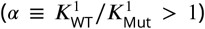 and increased resistance to drug 2 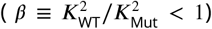, where superscripts label the drug (1 or 2) and subscripts label the cell type (wild type, WT; mutant, Mut).

**Figure 1.**
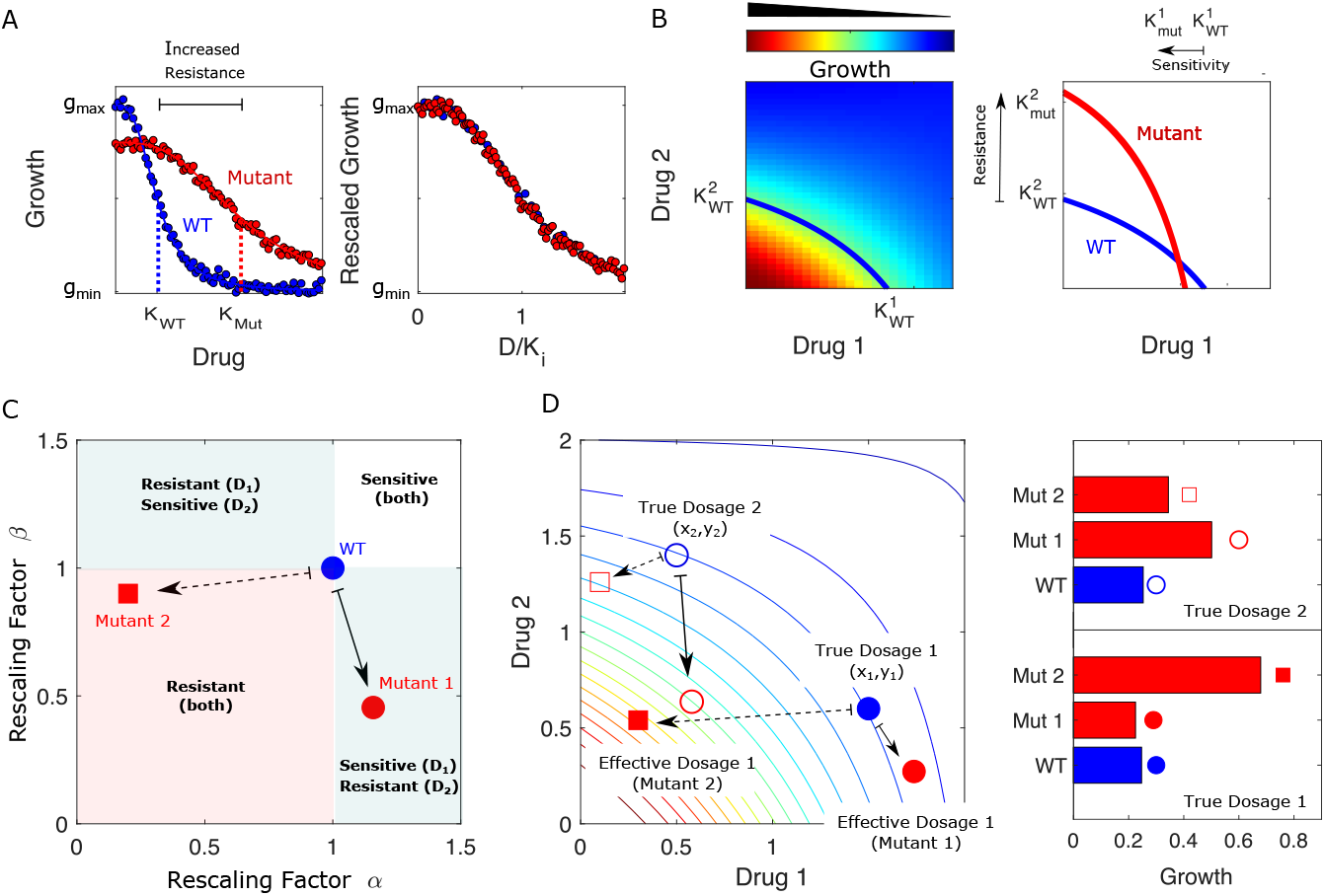
Drug resistance as a rescaling of effective drug concentration. The fundamental assumption of our model is that drug resistant mutants exhibit phenotypes identical to those of the ancestral (“wild type”) cells but at rescaled effective drug concentration. **A.** Left panel: schematic dose response curves for an ancestral strain (blue) and a resistant mutant (red). Half maximal inhibitory concentrations (*K*_WT_, *K*_Mut_), which provide a measure of resistance, correspond to the drug concentrations where inhibition is half maximal. Fitness cost of resistance is represented as a decrease in drug-free growth. Right panel: dose response curves for both cell types collapse onto a single functional form, similar to those in (***Chait et al., 2007***; ***Michel et al., 2008***; ***Wood and Cluzel, 2012***; ***Wood et al., 2014***). B. Left panel: in the presence of two drugs, growth is represented by a surface; the thick blue curve represents the isogrowth contour at half maximal inhibition; it intersects the axes at the half maximal inhibitory concentrations for each individual drug. Right panel: isogrowth contours for ancestral (WT) and mutant cells. In this example, the mutant exhibits increases resistance to drug 2 and an increased sensitivity to drug 1, each of which corresponds to a rescaling of drug concentration for that drug. These rescalings are quantified with scaling constants 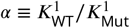 and 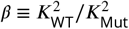, where the superscripts indicate the drug (1 or 2). **C.**Scaling factors for two different mutants (red square and red circle) are shown. The ancestral cells correspond to scaling constants *α = β* = 1. Mutant 1 exhibits increased sensitivity to drug 1 (*α* > 1) and increased resistance to drug 2 (*β* < 1). Mutant 2 exhibits increased resistance to both drugs (*α < β* < 1), with higher resistance to drug 1. **D.** Scaling parameters describe the relative change in effective drug concentration experienced by each mutant. While scaling parameters for a given mutant are fixed, the effects of those mutations on growth depend on the external environment (i.e. the drug dosage applied). This schematic shows the effective drug concentrations experienced by WT cells (blue circles) and the two different mutants (red circles and red squares) from panel C under two different external conditions (open and closed shapes). True dosage 1 (2) corresponds to higher external concentrations of drug 1 (2). The concentrations are superimposed on a contour plot of the two drug surface (similar to panel B). Right panel: resulting growth of mutants and WT strains at dosage 1 (bottom) and dosage 2 (top). Because the dosages are chosen along a contour of constant growth, the WT exhibits the same growth at both dosages. However, the growth of the mutants depends on the dose, with mutant 1 growing faster (slower) than mutant 2 under dosage 2 (dosage 1). A key simplifying feature of these evolutionary dynamics is that the selective regime (drug concentration) and phenotype (effective drug concentration) have same units.

The power of this rescaling approach is that it directly links growth of the mutant populations to measurable properties of the ancestral population (the two-drug response surface) via traits of the mutants (the rescaling parameters). Each mutant is characterized by a pair of scaling parameters, (*α_i_, β_i_*), which one might think of as a type of coarse-grained genotype (Figure 1C). When paired with the ancestral growth surface, these traits fully determine the per capita growth rate of the mutant at any external dosage combination (*x, y*) via Equation 1. While the scaling parameters are intrinsic properties of each mutant, they contribute to the phenotype (growth) in a context-dependent manner, leading to selection dynamics that depend in predictable ways on the external environment (Figure 1).

We assume a finite set of *M* subpopulations (mutants), (*i* = 1, ..*M*), with each subpopulation corresponding to a single pair of scaling parameters. For simplicity, we assume each of these mutants is initially present in the population at low frequency and neglect mutations that could give rise to new phenotypes, though it is straightforward to incorporate them into the same framework. We do not specify the mechanisms by which this standing variation is initially generated or maintained; instead, we simply assume that such standing variation exists, and our goal is to predict how selection will act on this variation for different choices of external (true) drug concentrations. As we will see, statistical properties of this variation combine with the local geometry of the response surface to determine the selection dynamics of these traits.

### Population dynamics of scaling parameters

The mean scaling parameter 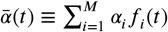 evolves according to

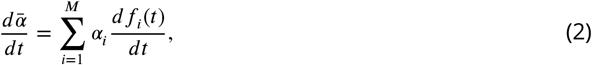

where *f_i_*(*t*) is the population frequency of mutant *i* at time *t* in the population. Assuming that each subpopulation grows exponentially at the per capita growth rate (*dn_i_/dt = g_i_n_i_*, with *n_i_* the abundance of mutant *i* and *g_i_* the per capita growth rate given by Equation 1), the frequency *f_i_*(*t*) changes according to

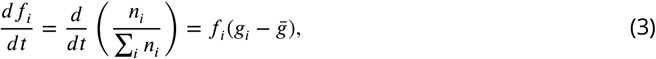

where 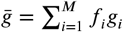 is the (time-dependent) mean value of *g_i_* across all *M* subpopulations (mutants). Combining Equations 2-3, we have

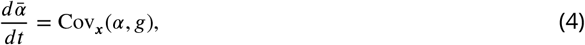

where 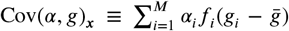 is the covariance between the scaling parameters *α_i_* and the corresponding mutant growth rates *g_i_*. The subscript ***x*** is a reminder that the growth rates *g_i_* and 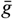 that appear in the covariance sum depend on the external (true) drug concentration ***x*** ≡ (*x, y*). An identical derivation leads to an analogous equation for the scaling parameter relative to drug 2, 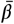, and the full dynamics are therefore described by

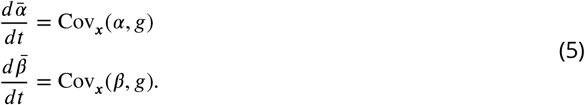

To complete the model described by Equation 5, one must specify the external (true) concentration of each drug (***x***); a finite set of scaling parameter pairs *α_i_, β_i_* corresponding to all “available” mutations; and an initial condition 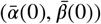 for the mean scaling parameters. When combined with the external drug concentrations, the scaling parameters directly determine the effective drug concentrations 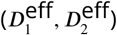 experienced by each mutant according to

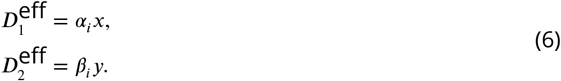

We note that drug concentrations can be above or below the minimum inhibitory concentration (MIC) contour, with higher concentrations leading to population collapse (*G* < 0) and lower concentrations to population growth (*G* > 0) in the ancestral (drug sensitive) cells. However selection dynamics remain the same in both cases, as selection depends only on differences in growth rates between different subpopulations.

Equation 5 is an example of the well-known Price Equation from evolutionary biology (***Price, 1970, 1972***; ***Frank, 1995***; ***Lehtonen et al., 2020***), which says that the change in the (population) mean value of a trait is governed by the covariance of traits and fitness. In general, fitness can be difficult to measure and, in some cases, even difficult to define. However, the rescaling assumption of our model replaces fitness with *g*, which can be directly inferred from measurable properties (the two drug response surface) in the ancestral population.

Equation 5 encodes deceptively rich dynamics that depend on both the interaction between drugs and the collateral effects of resistance. First, it is important to note that *α*’s and *β*’s vary together in pairs, and the evolution of these two traits is not independent. As a result, constraints on the joint co-occurrence of *α_i_* and *β_i_* among the mutant sub-populations can significantly impact the dynamics. These constraints correspond to correlations between resistance levels to different drugs-that is, to cross resistance (when pairs of scaling parameters simultaneously increase resistance to both drugs) or to collateral sensitivity (when one scaling parameter leads to increased resistance and the other to increased sensitivity). In addition, *g* contains information about the dose response surface and, therefore, about the interaction between drugs. The evolution of the scaling parameters is not determined solely by the drug interaction or by the collateral effects, but instead by both-quantified by the covariance between these rescaled trait values and the ancestral dose response surface.

As an example, we solved Equation 5 numerically to determine the dynamics of the mean scaling parameters and the mean growth rate for a population exposed to a fixed concentration of two drugs whose growth surface has been fully specified (Figure 2A). The dynamics can be thought of as motion on the two-dimensional response surface; if the initial population is dominated by the ancestral cells, the mean scaling parameters are approximately 1, and the trajectory therefore starts near the point representing the true drug concentration (in this case, (*x*_0_, *y*_0_)), where growth is typically small (Figure 2A). Over time, the mean traits evolve, tracing out a trajectory in the space of scaling parameters ((Figure 2B). When the external concentration of drug is specified, these dynamics also correspond to a trajectory through the space of effective drug concentrations which, in turn, can be linked with an average growth rate through the drug response surface (Figure 2B). The model therefore describes both the dynamics of the scaling parameters, which describe how resistance to each drug changes over time (Figure 2C), and the dynamics of growth rate adaptation in the population as a whole (Figure 2D).

**Figure 2.**
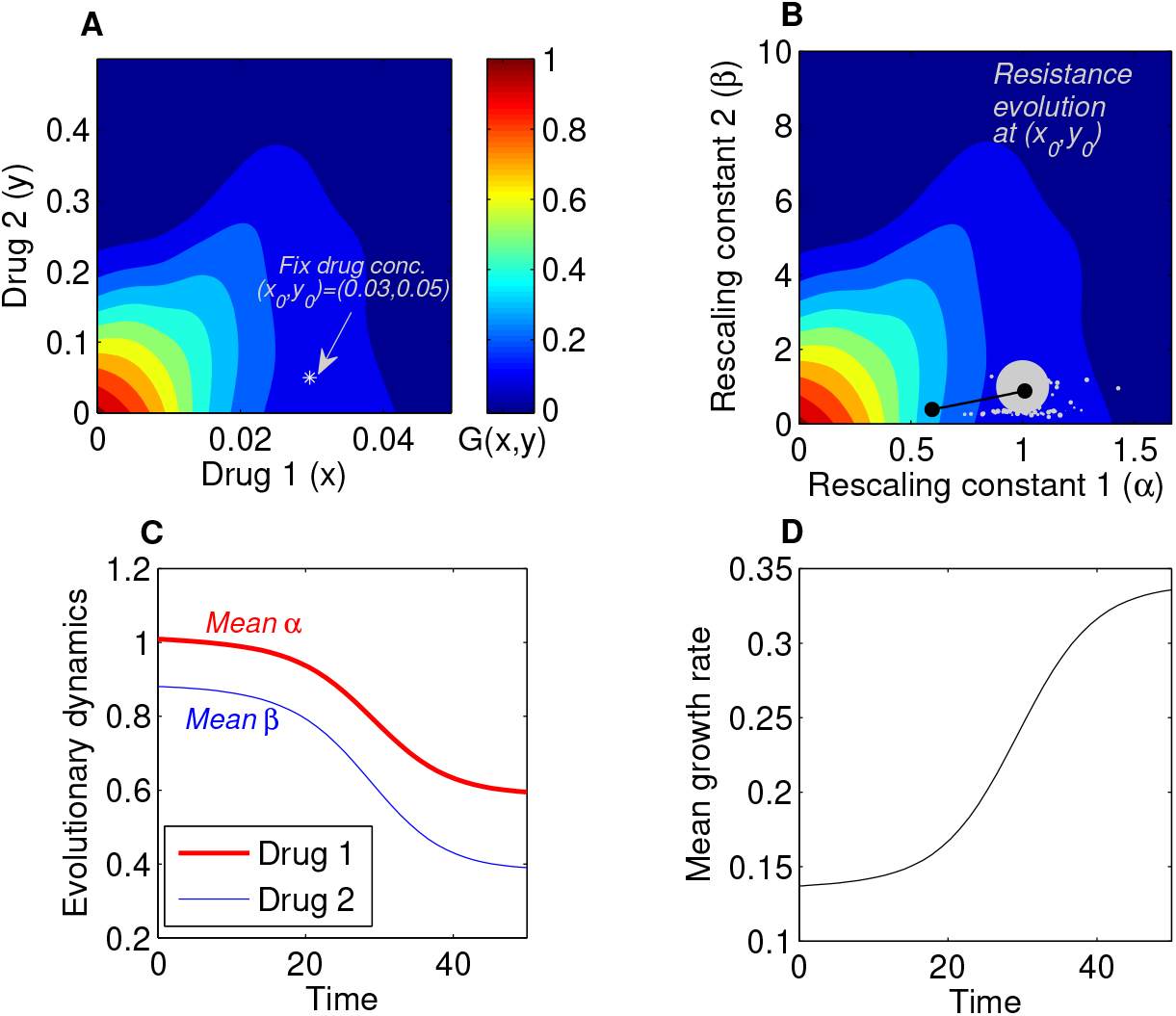
Using the Price Equation framework for predicting resistance evolution to two drugs in a population of bacterial cells. **A.** We used a growth landscape and available standing variation that were empirically derived (see Supplementary Figure S2-C). Drug 1 corresponds to Tigecycline and drug 2 corresponds to Ciprofloxacin. Bacterial population growth rate was determined at several combinations of concentrations of the two drugs (*x* ∈ [0, 0.05], *y* ∈ [0, 0.5]). We then simulated an external condition where the combination drug concentration is held constant at 0.6 of the maximal concentration of drug 1 and 0.1 of the maximal concentration of drug 2. In such environment, under a given random initial distribution of the population across susceptibility space, we predict the trajectory of evolution. **B.** The evolution of drug-resistance to two drugs in rescaling factor space. The WT is assumed to lie at (1,1) in the rescaling space and occupy 99 % of the total population. The available mutants, constituting the remaining 1%, are characterized by combinations of (*α_i_, β_i_*) traits and assigned random initial frequencies. **C**. The 2-dimensional trajectory can be decomposed in two time series of mean trait evolution where the mean traits reflect the rescaling constants with respect to the drug concentration applied 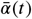 (red curve) and 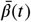 (blue curve). For the detailed selection dynamics see Supplementary Movie 1. **D.** The global mean fitness of the population during evolution is expected to increase, and can be computed from the selection dynamics solving the Price equation at each time-step.

### Selection dynamics depend on drug interaction and collateral effects

This rescaling model indicates that selection is determined by both the drug interaction and the collateral effects, consistent with previous experimental findings. As a simple example, consider the case of a fixed drug interaction but different qualitative types of collateral effects-that is, different statistical relationships between resistance to drug 1 (via *α*) and resistance to drug 2 (via *β*). In Figure 3A, we consider cases where resistance is primarily to drug 2 (black), primarily to drug 1 (cyan), strongly correlated (cross resistance, magenta), and strongly anticorrelated (collateral sensitivity, green). Using the same drug interaction surface as in Figure 2, we find that a mixture of both drugs leads to significantly different trajectories in the space of scaling parameters (Figure 3B) and, in turn, significantly different rates of growth adaptation. In this example, cross resistance (magenta) leads to rapid increases in resistance to both drugs (rapid decreases in scaling parameters, Figure 3C-D) and the fastest growth adaptation (Figure 3E). By contrast, if resistance is limited primarily to one drug (cyan or black), growth adaptation is slowed (Figure 3E)-intuitively, purely horizontal or purely vertical motion in the space of scaling parameters leads to only a modest change in growth because the underlying response surface is relatively flat in those directions (Figure 3).

**Figure 3.**
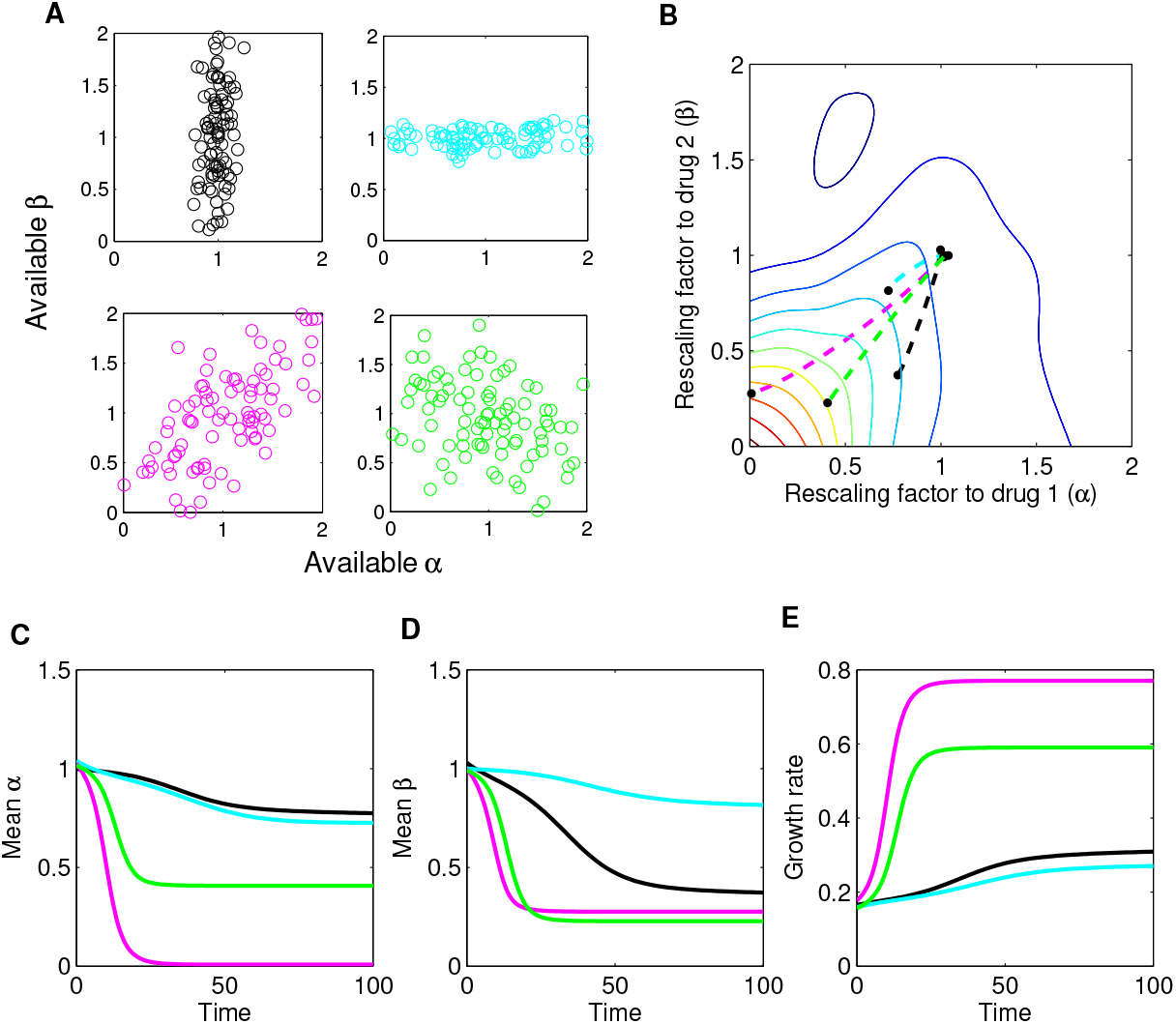
Different collateral profiles drive different evolutionary dynamics under the same treatment. Here, we simulate random collateral profiles for susceptibilities to two drugs and use them to predict phenotypic trajectories of a bacterial population. We assumed the growth landscape from Figure S2-C. **A.** Four special profiles include variation only in *β* (black), variation only in *α* (cyan), positive correlation between *α*’s and *β*’s (purple) and negative correlation between the two susceptibilites (green). **B.** The trajectories in mean *α − β* space following treatment (*x*_0_, *y*_0_), with *x*_0_ = 0.5*x_max_* and *y*_0_ = 0.5*y_max_*, corresponding to different collateral profiles. **C** The dynamics of mean susceptibility to drug 1 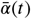 for the four cases. **D** The dynamics of mean susceptibility to drug 2 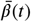 for the four cases. **E** The dynamics of mean growth rate for the four cases. For underlying heterogeneity, we drew 100 random *α_i_* and *β_i_* as shown in A, and initialized dynamics at ancestor frequency 0.99 and the remaining 1% evenly distributed among available mutants. It is clear from these graphs that the fastest increase in resistance to two drugs and increase in growth rate occurs for the collateral resistance (positive correlation) case. The time course of the detailed selective dynamics in these four cases are depicted in Supplementary Movies 2-5.

When both collateral effects and drug interactions vary, the dynamics can be considerably more complex, and the dominant driver of adaptation can be drug interaction, collateral effects, or a combination of both. Previous studies support this picture, as adaptation has been observed to be driven primarily by drug interactions (***Chait et al., 2007***; ***Michel et al., 2008***; ***Hegreness et al., 2008***), primarily by collateral effects (***Munck et al., 2014***; ***Barbosa et al., 2018***), or by combinations of both (***Baym et al., 2016b***; ***Barbosa et al., 2018***; ***Dean et al., 2020***). Figure 4 shows schematic examples of growth rate adaptation for different types of collateral effects (rows, ranging from cross resistance (top) to collateral sensitivity (bottom)) and drug interactions (columns, ranging from synergy (left) to antagonism (right)). The growth adaptation may also depend sensitively on the external environment-that is, on the true external drug concentration (blue, cyan, and red). In the absence of drug interactions (linear isoboles) and collateral effects (uncorrelated scaling parameters), adaptation is slower when the drugs are used in combination than when they are used alone, consistent with the fact that adaptation to multiple stressors is expected to take longer than adaptation to each stressor alone (Figure 4; middle row, middle column). As in Figure 3, modulating collateral effects with drug interaction fixed can have dramatic impacts on adaptation (Figure 4, columns). On the other hand, modulating the drug interaction in the absence of collateral effects will also significantly impact adaptation, with synergistic interactions leading to accelerated adaptation relative to both 1) other types of drug interaction (Figure 4, middle row; compare green curves across row) and 2) single drug adaptation (Figure 4, middle row, left panel). Similar interaction-driven adaptation has been observed in multiple bacterial species (***Chait et al., 2007***; ***Michel et al., 2008***; ***Hegreness et al., 2008***; ***Dean et al., 2020***).

**Figure 4.**
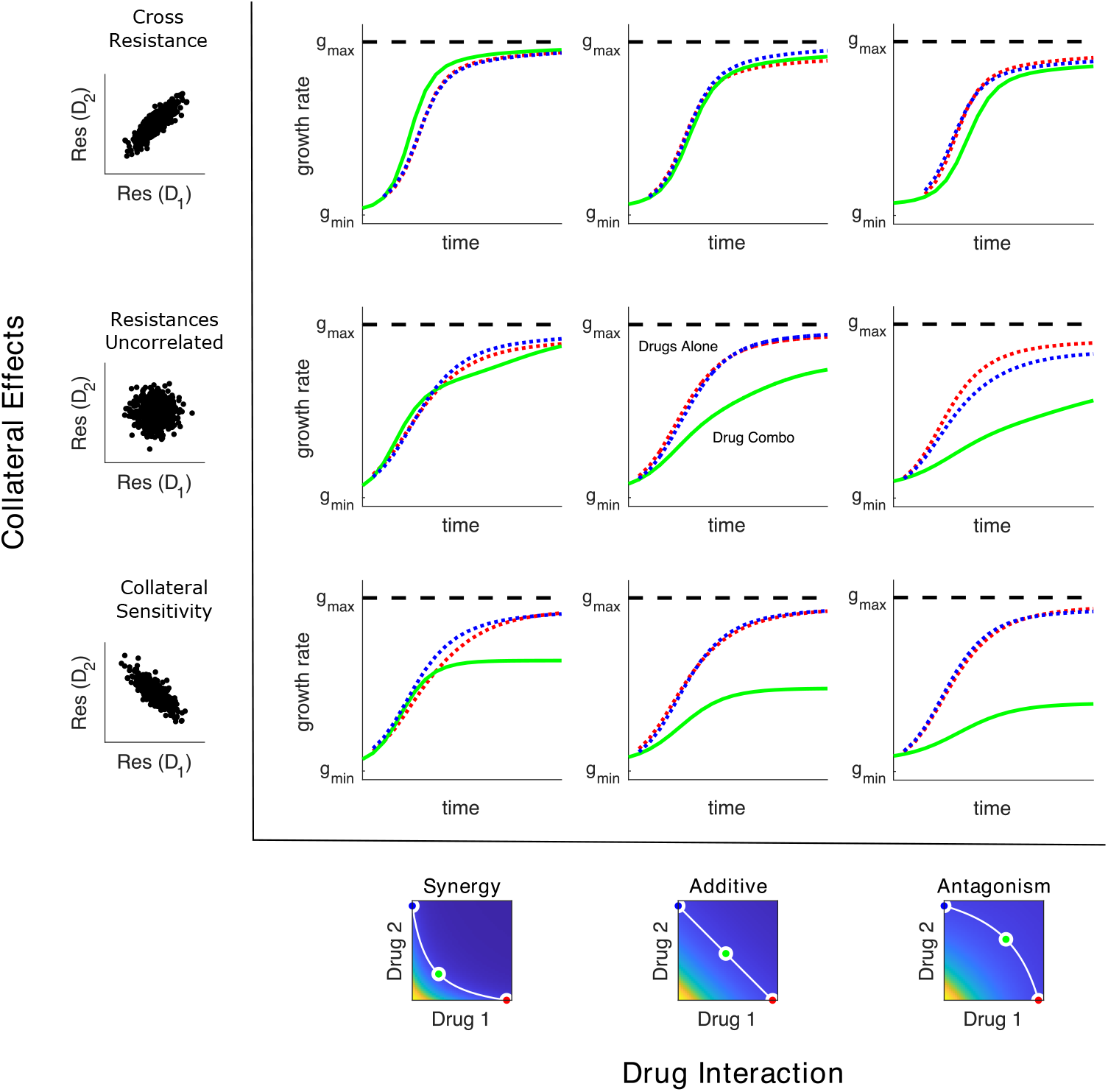
Adaptation depends on drug interaction and collateral effects of resistance. Drug interaction (columns) and collateral effects (rows) modulate the rate of growth adaptation (main 9 panels). Evolution takes place at one of three dosage combinations (blue, drug 2 only; cyan, drug combination; red, drug 1 only) along a contour of constant growth in the ancestral growth response surface (bottom panels). Drug interactions are characterized as synergistic (left), additive (center), or antagonistic (right) based on the curvature of the iso-growth contours. Collateral effects describe the relationship between the resistance to drug 1 and the resistance to drug 2 in an ensemble of potential mutants. These resistance levels can be correlated (top row, leading to collateral resistance), uncorrelated (center row), or anti-correlated (third row, leading to collateral sensitivity). Growth adaptation (main 9 panels) is characterized by growth rate over time, with dashed lines representing evolution in single drugs and solid lines indicating evolution in the drug combination. In this example, response surfaces are generated with a classical pharmacodynamic model that extends Loewe additivity by including a single drug interaction index that can be tuned to create surfaces with different combination indices (***Greco et al., 1995***).

### Selection as weighted gradient dynamics on the ancestral response surface

To gain intuition about the dynamics in the presence of both collateral effects and drug interactions, we consider an approximately monomorphic population where scaling parameters are initially narrowly distributed around their mean values. In this case, we can Taylor expand the function *g_i_* = *G*(*α_i_x*_0_, *β_i_y*_0_) (corresponding to the growth of mutant i) around the mean values 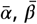, leading to

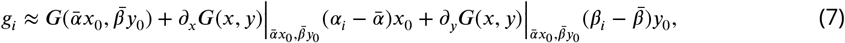

where we have neglected terms quadratic and higher. In this regime, *g_i_* is a linear function of the scaling parameters, and the covariances can therefore be written as

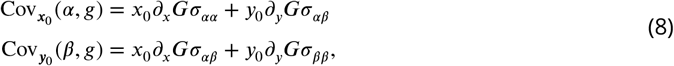

where 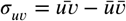 and we have used the fact that 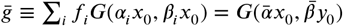 to first order. This is a type of weak-selection approximation: the trait that is evolving may have a very strong effect on fitness, but if there is only minor variation in such trait in the population, there will only be minor differences in fitness. Equation 5 therefore reduces to

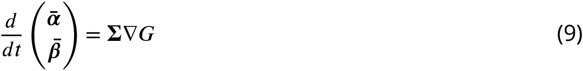

where 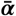 is a vector with components 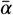 and 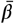, **∑** is a covariance-like matrix given by

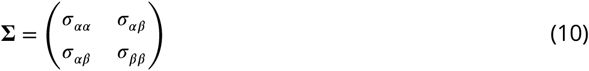

and Δ*G* is a weighted gradient of the function *G*(*x, y*) evaluated at the mean scaling parameters,

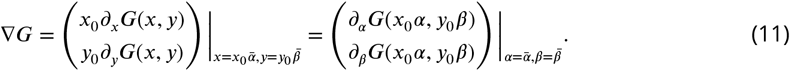

Equation 9 provides an accurate description of the full dynamics across a wide range of conditions (Figure S1) and has a surprisingly simple interpretation. Adaptation dynamics are driven by a type of weighted gradient dynamics on the response surface, with the weighting determined by the correlation coefficients describing resistance levels to the two drugs. In the absence of collateral effects-when **∑** is diagonal-the scaling parameters trace out trajectories of steepest ascent on the two-drug response surface. That is, in the absence of constraints on the available scaling parameters, adaptation follows the gradient of the response surface to rapidly achieve increased fitness, and because the response surface defines the type of drug interaction, that interaction governs the rate of adaptation. On the other hand, collateral effects introduce off-diagonal elements of **∑**, biasing trajectories away from pure gradient dynamics to account for the constraints of collateral evolution.

### Model predicts experimentally observed adaptation of growth and resistance

Our model makes testable predictions for adaptation dynamics of both the population growth rate and the population-averaged resistance levels to each drug (i.e. the mean scaling parameters) for a given drug-response surface, a given set of available mutants, and a specific combination of (external) drug dosages. To compare predictions of the model with experiment, we solved Equation 5 for the eleven different dosage combinations of Tigecycline (TGC) and Ciprofloxacin (CIP) used to select drug-resistance mutants in (***Dean et al., 2020***). We assumed that the initial population was dominated by ancestral cells (*α = β* = 1) but also included a subpopulation of resistant mutants whose scaling parameters were uniformly sampled from those measured across multiple days of the evolution (see Methods). The model predicts different trajectories for different external doses (selective regimes: red to blue, Figure 5, top panel), leading to dramatically different changes in resistance (IC_50_) and growth rate adaptation (Figure 5, bottom panels), similar to those observed in experiment. Specifically, the model predicts (and experiments confirm) a dramatic decrease in CIP resistance as TGC concentration is increased along a contour of constant growth. As TGC eclipses of critical concentration of approximately 0.025, selection for both CIP resistance and increased growth are eliminated. Intuitively, these dynamics result from the strongly antagonistic interactions between the two drugs, which reverses the selection for resistance to CIP; similar results have been seen in other species and for other drug combinations (***Chait et al., 2007***). We also compared model predictions with experimental adaptation to two additional drug pairs (Figures S3 and S4) and again found that the model captures the qualitative features of both resistance changes to the component drugs and growth adaptation.

**Table 1.** Different biological properties correspond to different features of the model

| Biological property | Model property |
| --- | --- |
| Scaling parameters (ancestral) | $\alpha = \beta = 1$ |
| Resistance to drug 1 (Relative MIC) | Inverse scaling parameter $\alpha^{-1}$ |
| Resistance to drug 2 (Relative MIC) | Inverse scaling parameter $\beta^{-1}$ |
| Synergistic interaction | Locally convex isoboles |
| Antagonistic interaction | Locally concave isoboles |
| Cross resistance | Correlated scaling parameters $(\alpha, \beta)$ |
| Collateral sensitivity | Anti-correlated scaling parameters $(\alpha, \beta)$ |
| External concentration $(x, y)$ | Effective drug concentration $(\alpha x, \beta y)$ |
| Fitness cost | Prefactor $g \rightarrow (1 - \gamma)g$ |

**Figure 5.**
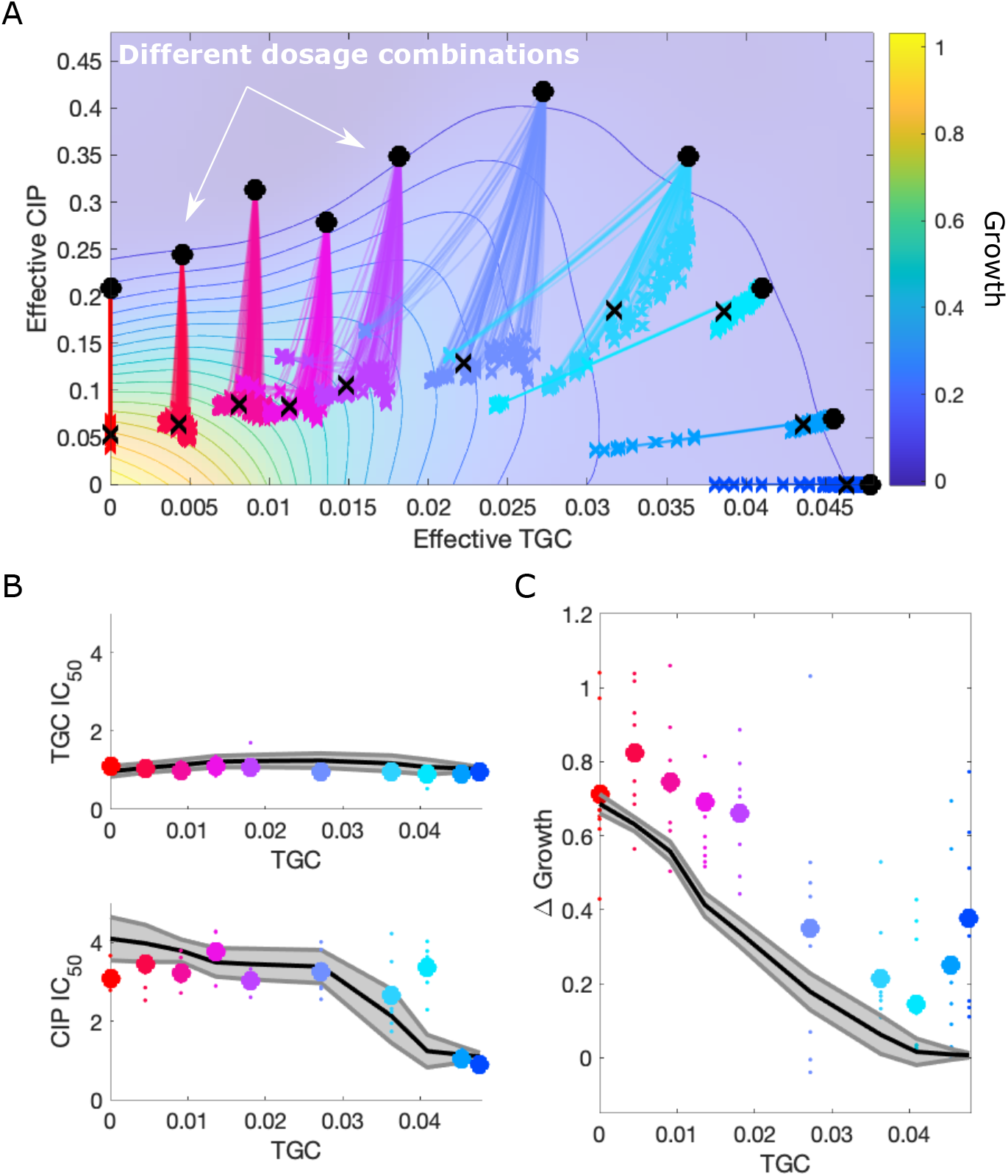
Price equation describes adaptation dynamics of growth and drug resistance in *E. faecalis* under different selective regimes. A. Experimentally measured growth response surface for tigecycline (TGC) and ciprofloxacin (CIP) from (***Dean et al., 2020***). Circles represent 11 different adaptation conditions, each corresponding to a specific dosage combination (*x*_0_, *y*_0_). Solid lines show adaptation trajectories (i.e. changes in effective drug concentration over time) predicted from the Price Equation. In each case, the set of available mutants–and hence, the set of possible scaling parameters *α* and *β*–is determined by sampling from experimentally measured changes in resistance (half-maximal inhibitory concentration IC_50_) for each drug in the full ensemble of mutants arising from adaptation to these drugs. B. Relative change in IC_50_ for populations adapted in each of the 11 conditions in the top panel. Each solid curve is a prediction from the Price Equation for one particular sampling of available mutants. Small circles are results from individual populations; larger markers are averages over all populations adapted to a given condition. C. Change in population growth rate for populations adapted in each condition. Each solid curve is a prediction from the Price Equation for one particular sampling of available mutants. Small circles are results from individual populations; larger markers are averages over all populations adapted to a given condition.

### Evolutionary dynamics under temporal sequences of drug combinations

Evolutionary dynamics in the presence of multiple drugs can be extremely complex. While past work has focused primarily on the use of either temporal sequences or (simultaneous) combinations of antibiotics, more complex scenarios are possible, in principle, but difficult to analyze even in theoretical models. The simplicity of our model allows us to investigate evolutionary dynamics in response to a more complex scenario: temporal *sequences* of antibiotic *combinations*. As proof-of-principle, we numerically studied time-dependent therapies consisting of two sequential regimes, deemed treatment A and treatment B (Fig. 6). Both regimes consist of a specific combination of two drugs, with the combinations chosen to lie along a contour of constant growth (Fig. 6A). As a result, the net inhibitory effect of each combination is the same, but because the drugs are given in different dosage ratios, the dynamics resulting from each treatment are expected to differ. Using experimental estimates of variability in *α, β* traits ((Fig. S2C), we find that resistance overall increases and growth rate reaches higher levels with treatment, but both the relative duration and the ordering between treatments matters. Strikingly we find that the order B+A (“order 2”) is superior to A+B (“order 1”) in constraining growth rate (Fig. 6J, M), even though the relative final mean resistance achieved is not particularly sensitive to the ordering (Fig. 6H, K; I, L). Furthermore, if we fix the total duration of treatment (Fig. 6O), we find that it is not only the proportion of time a certain drug-dosage combination is received that matters, but especially whether it is administered first or second. In this example, resistance evolves slowly and monotonically to drug 1, but more rapidly and non-montonically to drug 2, as the proportion of time in treatment A is tuned from 0 to 1 (Fig. 6P-Q). In particular, the model suggests two options for constraining resistance to drug 1: a short duration of A followed by a long duration of B, or a short duration of B followed by long duration of A (Fig. 6P). A similar pattern occurs for resistance to drug 2: order A+B is best if combined with short duration A, and order B+A is best if combined with short B (Fig. 6Q). The optimal way to constrain growth rate adaptation in this case is always A+B (order 1) with short A and long B. Interestingly, we observe that ordering effects do not matter for growth rate as the proportion of time in A approaches 1, but for resistance evolution it is precisely in the extremes of treatment partitioning that order effects matter more. In contrast, if A and B are equi-partitioned in time, then their order is less influential. This example illustrates how the complexity of the adaptive landscape and the underlying heterogeneity in antibiotic susceptibility profiles leads to rich and suprising dynamics. In principle, this model can be used to to identify cycles of antibiotic combinations that are more likely to manage resistance.

**Figure 6.**
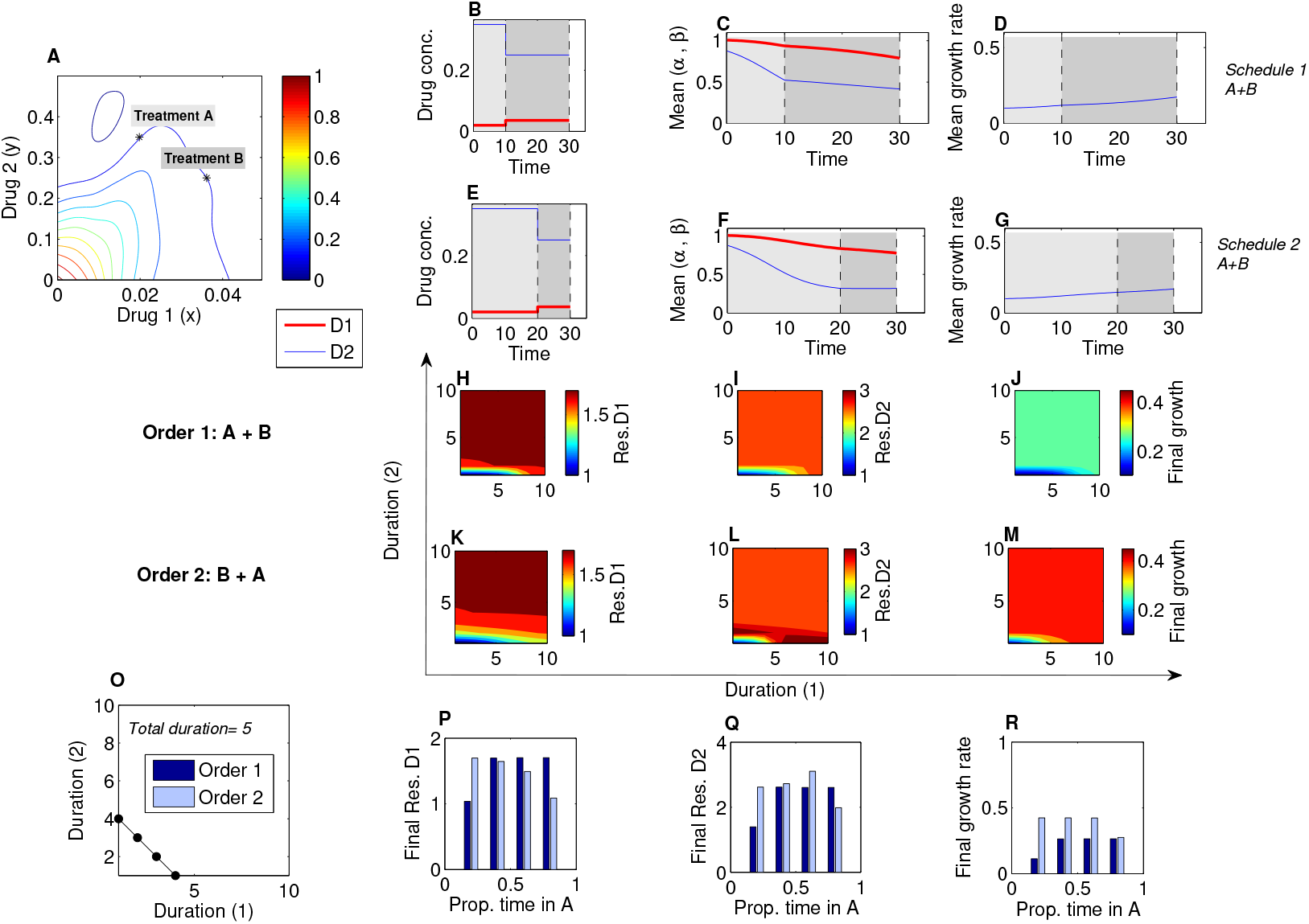
Sequential multi-drug treatment, under different schedules and treatment ordering. Here, we illustrate how the model framework can be used to predict phenotypic evolution and growth dynamics in a bacterial population under consecutive treatments with two drugs but administered at different dose combinations. For underlying heterogeneity, we assumed existing *α_i_* and *β_i_* as derived experimentally (Fig. S2C) and initialized dynamics at ancestor frequency 0.99 and the remaining 1% evenly distributed among available mutants. **A.** The growth landscape evaluated at different concentrations of two drugs. The marked asterisks denote two selected treatments (*x*_0_, *y*_0_) that are applied sequentially on this bacterial population. Treament A:*x*_0_=0.4*x_max_, y*_0_ = 0.7*y_max_* and Treatment B: *x*_0_ = 0.73*x_max_, y*_0_ = 0.5*y_max_*. **B-D.** Schedule 1 of short treatment A and longer treatment B. **E-G**Schedule 2 of longer treatment A and shorter treatment B. Different dynamics of 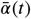 and 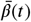 result from these schedules (**C,F**) and different adaptation in growth rate (**D,G**). **H-J** Final traits post-treatment 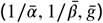 evolved under two multi-drug therapies (A+B) at different respective durations. **K-M** Final traits post-treatment evolved under two multi-drug therapies given in reverse order (B+A) at different respective durations. **O** Different ways of administering two consecutive combination therapies of a fixed total duration, here assumed 5. **P-R** Comparison of ordering effects when the proportion of treatment A is held fixed. Final traits at the end of the same therapeutic length, evolved under two multi-drug therapies given in order 1 and 2 (A first, vs. B first).

## Discussion

Antibiotic resistance is a growing threat to modern medicine and public health. Multi-drug therapies are a promising approach to both increase efficacy of treatment and prevent evolution of resistance, but the two effects can be coupled in counter-intuitive ways through the drug interactions and collateral effects linking the component drugs. Our results provide a unified model for incorporating both drug interactions and collateral effects to predict phenotypic adaptation on coarse-grained fitness landscapes that are measurable features of the ancestral population. Special cases of the model reproduce previous experimental results that appear, on the surface, to be contradictory; indeed, adaptation can be driven primarily by drug interactions, primarily by collateral effects, or by a combination of both, and the balance of these effects can be shifted using tunable properties of the system (i.e. the ratio of drug dosages). Our model was inspired by rescaling arguments that were originally introduced in (***Chait et al., 2007***) and have since been shown to capture phenotypic features of dose response surfaces or adaptation in multiple bacterial species (***Michel et al., 2008***; ***Hegreness et al., 2008***; ***Torella et al., 2010***; ***Wood and Cluzel,2012***; ***Wood et al., 2014***; ***Dean et al., 2020***; ***Das et al., 2020***). Our results complement these studies by showing how similar rescaling assumptions, when formalized in a population dynamics model, lead to testable predictions for the dynamics of both growth adaptation and phenotype (resistance) evolution. Importantly, the model also has a simple intuitive explanation, with evolutionary trajectories driven by weighted gradient dynamics on two-dimensional landscapes, where the local geometry of the landscape reflects the drug interaction and collateral effects constrain the direction of motion.

It is important to keep in mind several limitations of our approach. The primary rescaling assumption of the model is that growth of a drug resistant mutant is well approximated by growth of the ancestral strain at a new “effective” drug concentration-one that differs from the true external concentration. This approximation has considerable empirical support (***Chait et al., 2007***; ***Wood and Cluzel, 2012***; ***Wood et al., 2014***; ***Das et al., 2020***) but is not expected to always hold; indeed, there are examples where mutations lead to more complex nonlinear transformations of the drug response surface (***Wood et al., 2014***; ***Munck et al., 2014***). While these effects could in principle be incorporated into our model-for example, by assuming transformations of the ancestral surface, perhaps occurring on a slower timescale, that go beyond simple rescaling-we have not focused on those cases. For simplicity, we have also neglected a number of features that may impact microbial evolution. For example, we have assumed that different subpopulations grow exponentially, neglecting potential interactions including clonal interference (***Gerrish and Lenski, 1998***), intercellular (***Koch et al., 2014***; ***Hansen et al., 2017, 2020***) or intra-lineage (***Ogbunugafor and Eppstein, 2016***) competition, and cooperation (***Yurtsev et al., 2013***; ***Sorg et al., 2016***; ***Estrela and Brown, 2018***; ***Frost et al., 2018***; ***Hallinen et al., 2020***), though these complexities could also be incorporated in our model, perhaps at the expense of some intuitive interpretations (e.g. weighted gradient dynamics) that currently apply. We have also neglected *de novo* mutation and instead focused on selection driven by standing variation in the initial population, though as with the classic Price Equation, mutational effects-including stress-dependent modulation of mutation rate (***Kohanski et al., 2010***; ***Vasse et al., 2020***)– may be included as an additional term in Equation 5. In addition, we have not explicitly included a fitness cost of resistance (***Andersson and Hughes, 2010***)–that is, we assume that growth rates of mutants and ancestral cells are identical in the absence of drug. This assumption could be relaxed by including a prefactor to the growth function, *g*, → (1 – *γ_i_*(*α_i_, β_i_*))*g_i_*, where *γ_i_*(*α_i_, β_i_*) is the cost of resistance, which in general depends on the scaling parameters (if not, it can be easily incorporated as a constant). While such fitness costs have traditionally been seen as essential for reversing resistance (with, for example, drug-free “holidays” (***Dunai et al., 2019***)), our results underscore the idea that reversing adaptation relies on differential fitness along a multi-dimensional continuum of environments, not merely binary (drug / no drug) conditions. Our results indicate resistance and bacterial growth can be significantly constrained by optimal tuning of multi-drug environments, even in the absence of fitness cost. Finally, our model deals only with heritable resistance and therefore may not capture phenotypic affects associated with, for example, transient resistance (***El Meouche and Dunlop, 2018***) or cellular hysteresis (***Roemhild et al., 2018***).

One advantage of our approach is that it connects evolutionary effects of drug combinations and collateral effects with well established concepts in evolutionary biology and population genetics (***Price, 1970, 1972***; ***Day and Gandon, 2006***). Approximations similar to Equation 9 have been derived previously in quantitative genetics models (***Abrams et al., 1993***; ***Taylor, 1996***) and other applications of the Price Equation (***Lehtonen, 2018***). While the gradient approximation does not require that the population be monomorphic with rare mutants or a particular form for the phenotype distribution, it does require that the majority of trait values (here scaling parameters) are contained in a regime where *G*(*x, y*) is approximately linear; in our case, that linearity arises by Taylor expansion and neglecting higher-order deviations from the population mean. More generally, the direction of evolutionary change in our model is determined by the gradient of the fitness function ▽*G* evaluated at the mean trait 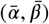; when the gradient vanishes, this point corresponds to a singular point (***Waxman and Gavrilets, 2005***) or an evolutionarily singular strategy (***Geritz et al., 1998***). Whether such point may be reached (convergence stability), and how much variance in trait values can be maintained around such point (whether evolutionarily stable, ESS), depend on other features, such as higher-order derivatives (***Otto and Day, 2011***; ***Eshel et al., 1997***; ***Lehtonen, 2018***; ***Smith, 1982***; ***Parker and Smith, 1990***). In the case of drug interactions, the fitness landscape in (*α, β*) space will typically have a single maximum at (0,0) corresponding to effective drug concentrations of zero. However, whether that point is reachable in general, or within a given time frame, will depend on the available mutants pre-existing at very low frequencies in the population, or on the speed and biases in the mutational process itself, if such mutants are to be generated *de novo* during treatment. In principle, the model also allows for long-term coexistence between different strains; in that case, the rescaled effective drug concentrations experienced by both strains would fall along a single contour of constant growth. Hence, while variance in the population growth will necessarily decrease over time, variance in the traits (scaling parameters) themselves can change non-monotonically.

The model can be extended in straightforward ways to incorporate additional details. For example, prescribing specific temporal or statistical constraints on the set of available scaling parameters could be used to incorporate information about rugged features of the underlying genetic landscapes, leading to historical contingencies between sequentially acquired mutations (***Barbosa et al., 2017***; ***Card et al., 2019***; ***Das et al., 2020***). The framework is sufficiently flexible to integrate different strands of empirical data, and our results underscore the need for quantitative phenotype data tracking resistance to multiple drugs simultaneously, especially when drug combinations are potentially driving selection dynamics. At an epidemiological level, the dominant approach in describing resistance has been to use fixed break-points in MIC and track the percentage of isolates with MIC above (drug-resistant) or below (drug-sensitive) such break-point (e.g.(***ECDC, 2019***; ***Chang et al., 2015***)). By missing or decoupling patterns of co-occurrence between MICs to different drugs across isolates, this approach remains incomplete for mechanistic predictions. Our framework suggests that going beyond such binary description towards a more continuous and multidimensional phenotype characterization of drug resistance is possible, with applications not just in microbiology but also in the evolutionary epidemiology of drug-resistance (***Day and Gandon, 2012***; ***Day et al., 2020***). In the long run, these advances may yield better and more precise predictions of resistance evolution at multiple scales, and, in turn, optimized treatments that balance short term inhibitory effects of a drug cocktail with its inseparable, longer-term evolutionary consequences.

Perhaps most importantly, our approach provides a low-dimensional approximation to the high-dimensional dynamics governing the evolution of resistance. In contrast to the classical genotype-centric approaches to resistance, our model uses rescaling arguments to connect measurable traits of resistant cells (scaling parameters) to environment-dependent phenotypes (growth). This rescaling dramatically reduces the complexity of the problem, as the two-drug response surfaces-and, effectively, the fitness landscape–can be estimated from only single drug dose response curves. Such coarse-grained models can help extract simplifying principles from otherwise intractable complexity (***Shoval et al., 2012***; ***Hart et al., 2015***). In many ways, the classical Price equation performsa similar function, revealing links between trait-fitness covariance and selection that, at a mathematical level, are already embedded in simple models of population growth. In the case of multi-drug resistance, this formalism reveals that drug interactions and collateral effects are not independent features of resistance evolution, and neither, alone, can provide a complete picture. Instead, they are coupled through local geometry of the two-drug response surface, and we show how specific dosage combinations can shift the weighting of these two effects, providing a framework for systematic optimization of time-dependent multi-drug therapies.

## Methods

### Estimating scaling parameters from experimental dose response curves

The scalingparametersfor a given mutant can be directly estimated by comparing single drug dose response curves of the mutant and ancestral populations. To do so, we estimate the half maximal inhibitory concentration (*K_i_*) for each population by fitting the normalized dose-response curve to a Hill-like function *g_i_*(*d*) = (1 + (*d/K_i_*)^*h*^)^−1^, with *g_i_*(*d*) the relative growth at concentration *d* and *h* a Hill coefficient measuring the steepness of the dose-response curve) using nonlinear least squares fitting. The scaling parameter for each drug is then estimated as the ratio of the *K_i_* parameters for the ancestral and mutant populations. For example, an increase in resistance corresponds to an increase in *K_i_* for the mutant population relative to that of the ancestor, yielding a scaling parameter of less than 1. Estimates for the scaling parameters for the 3 drug combinations used here are shown in Figure S2 (from data in (***Dean et al., 2020***)).

While it is straightforward to estimate the scaling parameters for any particular isolate, it is not clear *a priori* which isolates are present at time 0 of any given evolution experiment. To compare predictions of our model with lab evolution experiments, we first estimated scaling parameters for all isolates collected during lab evolution experiments in each drug pair (***Dean et al., 2020***). This ensemble includes 50-100 isolates per drug combination and includes isolates collected at different time points during the evolution (after days 1,2, or 3) as well as isolates selected in different dosage combinationsS2. We then randomly sampled from this ensemble to generate low-level standing diversity (on average, approximately 10 distinct pairs of scaling parameters) at time 0 for each simulation of the model, and we repeated this sub-sampling 100 times to generate an ensemble of evolutionary trajectories for each condition.

The results of the simulation can, in principle, depend on how these scaling parameters are sampled. While the qualitative differences between simulations do not depend heavily on this choice of sub-sampling in the data used here (Figure S5), one can imagine scenarios where details of the sub-sampling significantly impact the outcome. Similarly, precise comparison with experiment requires accurate estimates for the total evolutionary time and for the initial frequency of all resistant mutants, though for these data, the qualitative results do not depend sensitively on these choices (Figure S6 and S7). We stress that these are not fundamental limitations of the model itself, but instead arise because we do not have a precise measure of the standing variation characterizing these particular experiments. In principle, a more accurate ensemble of scaling parameters could be inferred from cleverly designed fluctuation tests (***Luria and Delbrück, 1943***) or, ideally, from high-throughput, single-cell phenotypic studies (***Baltekin et al., 2017***). At a theoretical level, subsampling could also be modulated to simulate the effects of different effective population sizes, with standing diversity expected to be significantly larger for large populations.

## Acknowledgements

This work was supported by Fundação Luso-Americana para o Desenvolvimento (FLAD/NSF grant-274/2016 to EG) and Instituto Gulbenkian de Ciência, the National Science Foundation (NSF No. 1553028 to KW), and the National Institutes of Health (NIH No. 1R35GM124875 to KW). The Center for Stochastic and Computational Mathematics is supported by FCT via UIDB/04621/2020 and UIDP/04621/2020. The format for this preprint is adapted from the eLife template available on Overleaf.com.

## Supplemental Figures and Movies

The supplemental information contains seven figures (S1-S7) and five movies (Movies 1-5).

### Movie 1

#### Detailed selection dynamics associated to Fig.2

Here we visualize the selection going on in a hypothetical bacterial population under the 2-drug dosage selective regime (*x*_0_, *y*_0_) and growth landscape shown in Figure 2, where the collateral effects, represented in the available mutants (*α_i_, β_i_*) are estimated from data (Fig.S2C). The size of the circles in rescaling factor trait space correspond to the relative frequency of each sub-population, calculated at each timepoint.

### Movie 2

#### Evolutionary dynamics for collateral effects in Fig.3A

Trajectory of mean traits in rescaling factor space (*α, β*) for case A. in Figure 3 where the randomly generated available standing variation is higher in *β* and much lower in *α*. The selective regime in drug dosage is (*x*_0_, *y*_0_) = 0.5(*x_max_, y_max_*). The size of the circles represents the relative frequencies of different mutant sub-populations available at the beginning.

### Movie 3

#### Evolutionary dynamics for collateral effects in Fig.3B

Trajectory of mean traits in rescaling factor space (*α, β*) for case A. in Figure 3 where the randomly generated available standing variation is lower in *β* and higher in *α*. The selective regime in drug dosage is (*x*_0_, *y*_0_) = 0.5(*x_max_, y_max_*). The size of the circles represents the relative frequencies of different mutant sub-populations available at the beginning.

### Movie 4

#### Evolutionary dynamics for collateral effects in Fig.3C

Trajectory of mean traits in rescaling factor space (*α, β*) for case A. in Figure 3 where the randomly generated available standing variation in *β* correlates positively with that in *α*. The selective regime in drug dosage is (*x*_0_, *y*_0_) = 0.5(*x_max_, y_max_*). The size of the circles represents the relative frequencies of different mutant sub-populations available at the beginning.

### Movie 5

#### Evolutionary dynamics for collateral effects in Fig.3B

Trajectory of mean traits in rescaling factor space (*α, β*) for case A. in Figure 3 where the randomly generated available standing variation in *β* correlates negatively with that in *α*. The selective regime in drug dosage is (*x*_0_, *y*_0_) =0.5(*x_max_, y_max_*). The size of the circles represents the relative frequencies of different mutant sub-populations available at the beginning.

**Figure S1.**
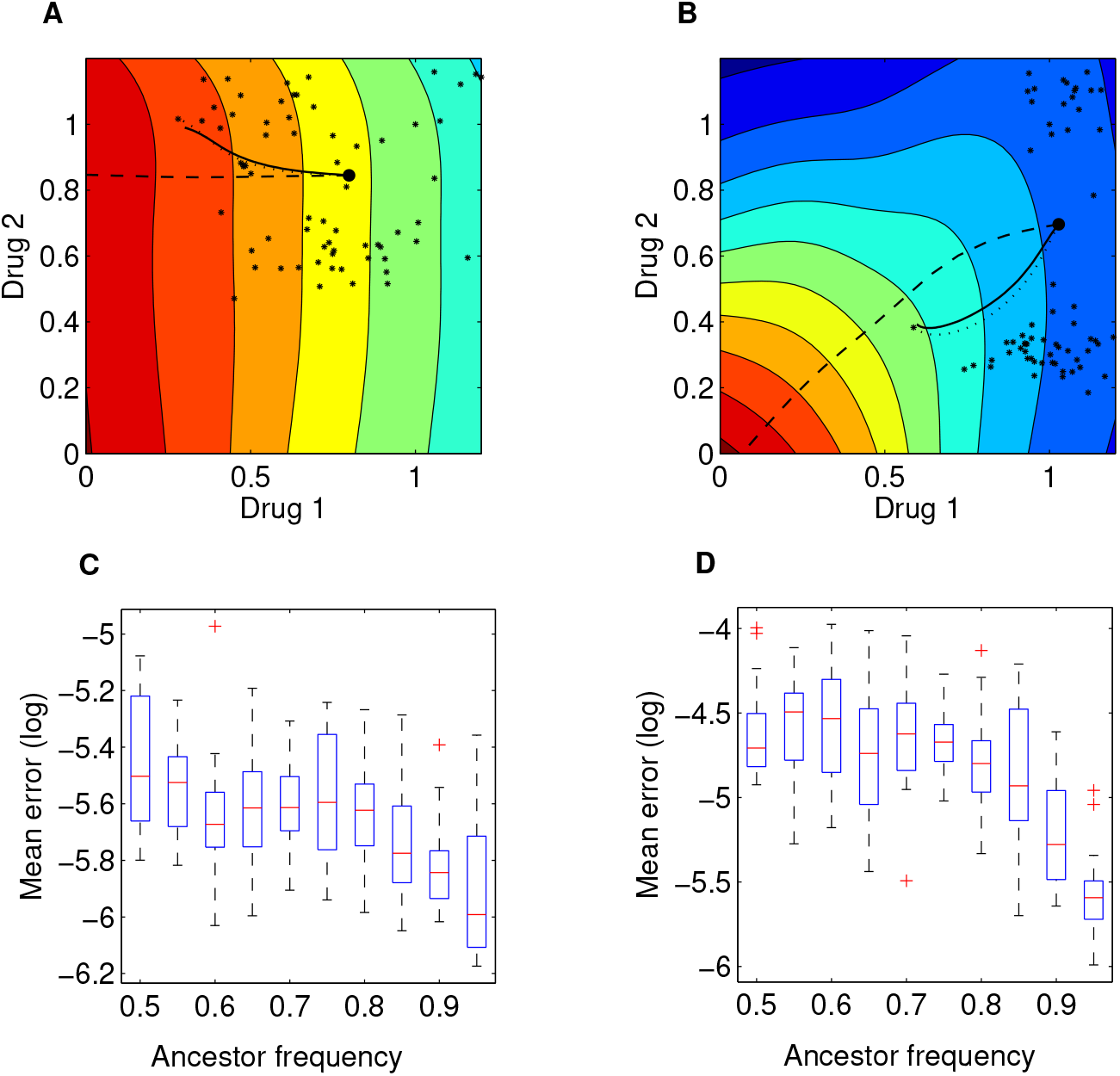
Illustration of the local growth gradient approximation to the Price equation. **A-B** Comparison between numerical solutions to the full Price Equation (dotted line), the local growth gradient approximation (solid line), and the trajectory of pure (unweighted) gradient descent (dashed line) for mean resistance evolution to two drugs 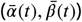. The dotted and solid lines are almost indistinguishable. In A and B, we assume two growth landscapes respectively (drug combination 1 vs. drug combination 3, see Supplementary Figure S2 A,C), external concentrations (*x*_0_, *y*_0_) = 0.4 × (*x_max_, y_max_*), and initial trait distributions following the empirically-derived (*α_i_, β_i_*). The large black dot represents the initial value 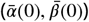, under initial ancestor frequency of 90% and the remaining 10% spread uniformly among available mutants (denoted via the small black asterisks in the 2-d trait space). **C-D**. Root mean square error between mean trait evolution calculated with the full iteration of the Price equation and the gradient approximation (for the two cases in A-B respectively), varying ancestor initial frequency. Here for each *p* ∈ (0.5,1) (the initial frequency of the ancestor sub-population with scaling parameters *α = β* = 1), we simulated 20 stochastic realizations of the evolutionary process under drug dosage (*x*_0_, *y*_0_), starting from different random frequencies for the remaining 1 – *p* proportion of the population composed of available mutants.

**Figure S2.**
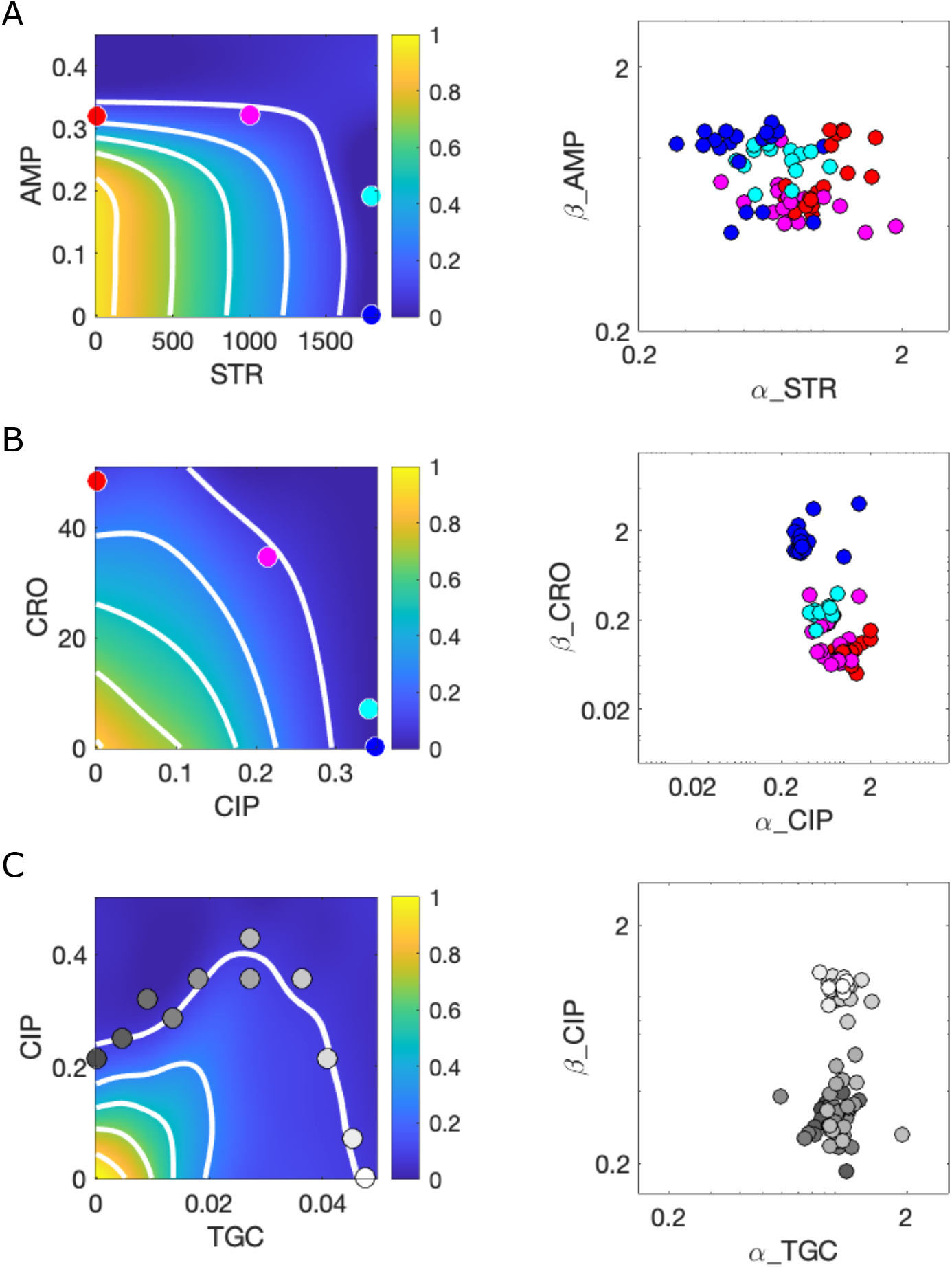
Response surfaces and scaling parameters for three drug pairs. Growth rate surfaces (left) and scaling parameters *α* and *β* for three drug combinations: ampicillin (AMP)-Streptomycin (STR) (**A**); ceftriaxone (CRO)-ciprofloxacin (CIP) (**B**); and ciprofloxacin (CIP)-tigecycline (TGC) (**C**). Large color-coded circles on the left panels indicate external concentrations at which selection took place. Circles on the right panels indicate all observed mutants (*α_i_, β_i_*) that emerged, pooled across conditions. Data from (***Dean et al., 2020***). Growth rate is measured in normalized units where growth of the drug-free ancestral cells (approximately 1 hr^−1^) is set to 1.

**Figure S3.**
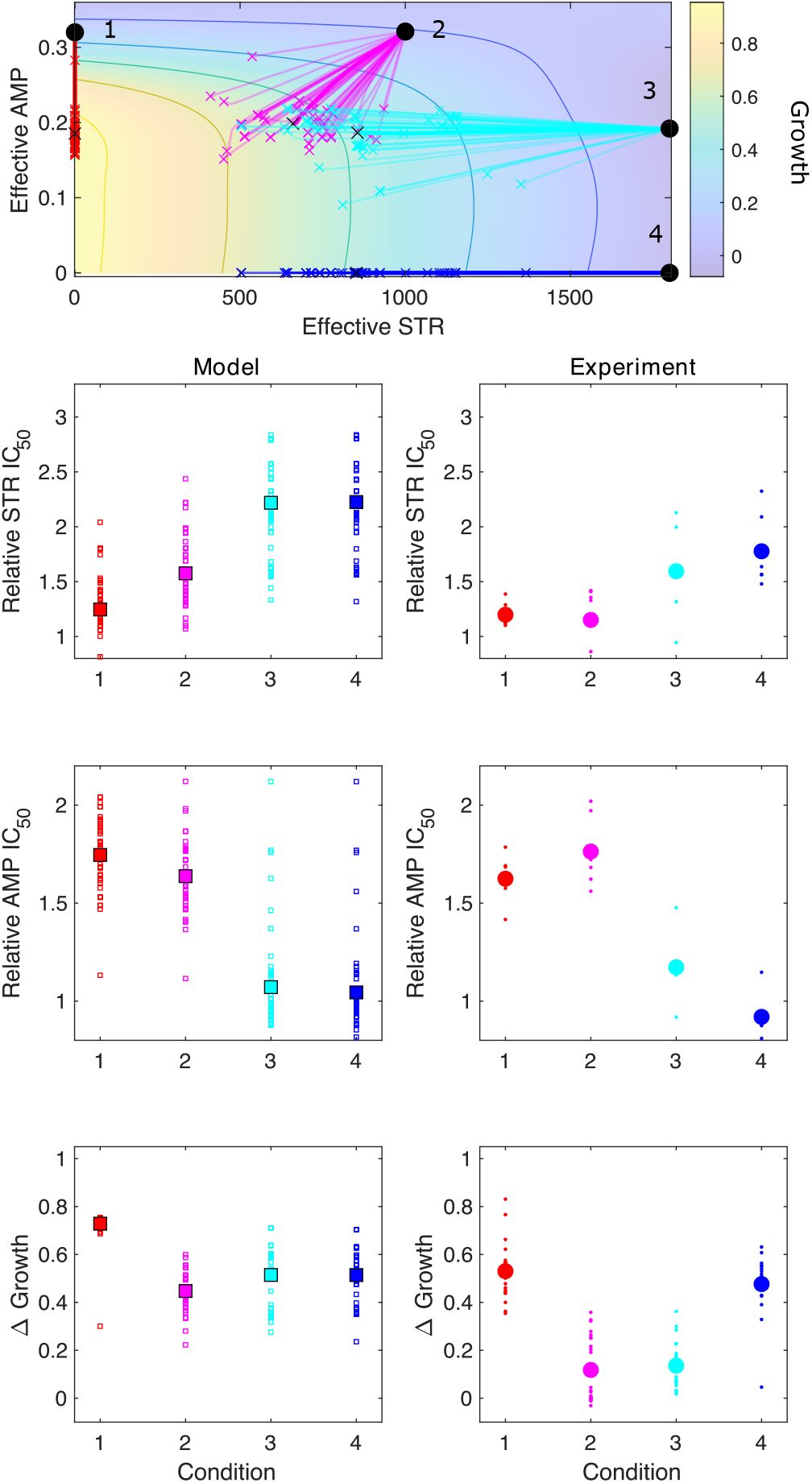
Model captures qualitative features of experimental evolution in combinations of AMP and STR. Top panel: experimentally measured growth response surface for ampicillin (AMP) and streptomycin (STR). Circles represent 4 different adaptation conditions, each corresponding to a specific dosage combination (*x*_0_, *y*_0_). Solid lines show adaptation trajectories (i.e. changes in effective drug concentration over time) predicted from the Price Equation over an interval t=48 hrs. In each case, the set of available mutants–and hence, the set of possible scaling parameters *α* and *β*-is determined by sampling from experimentally measured changes in resistance (half-maximal inhibitory concentration IC_50_) for each drug in the full ensemble of mutants arising from adaptation to these drugs (see Figure S2). Bottom panels: comparing results from the model (left column) and experiment (right panel). Quantities include relative change in IC_50_ for each drug (top two rows) as well as overall change in population growth (bottom row); in each case, results are shown for each of the 4 conditions in the top panel. For the experiments, small markers are results from individual populations;larger markers are averages over all populations adapted to a given condition. For the model, small markers are results from each sub-sampling of the experimental scaling parameters;larger markers are averages over all sub-samples. Experimental data from (***Dean et al., 2020***).

**Figure S4.**
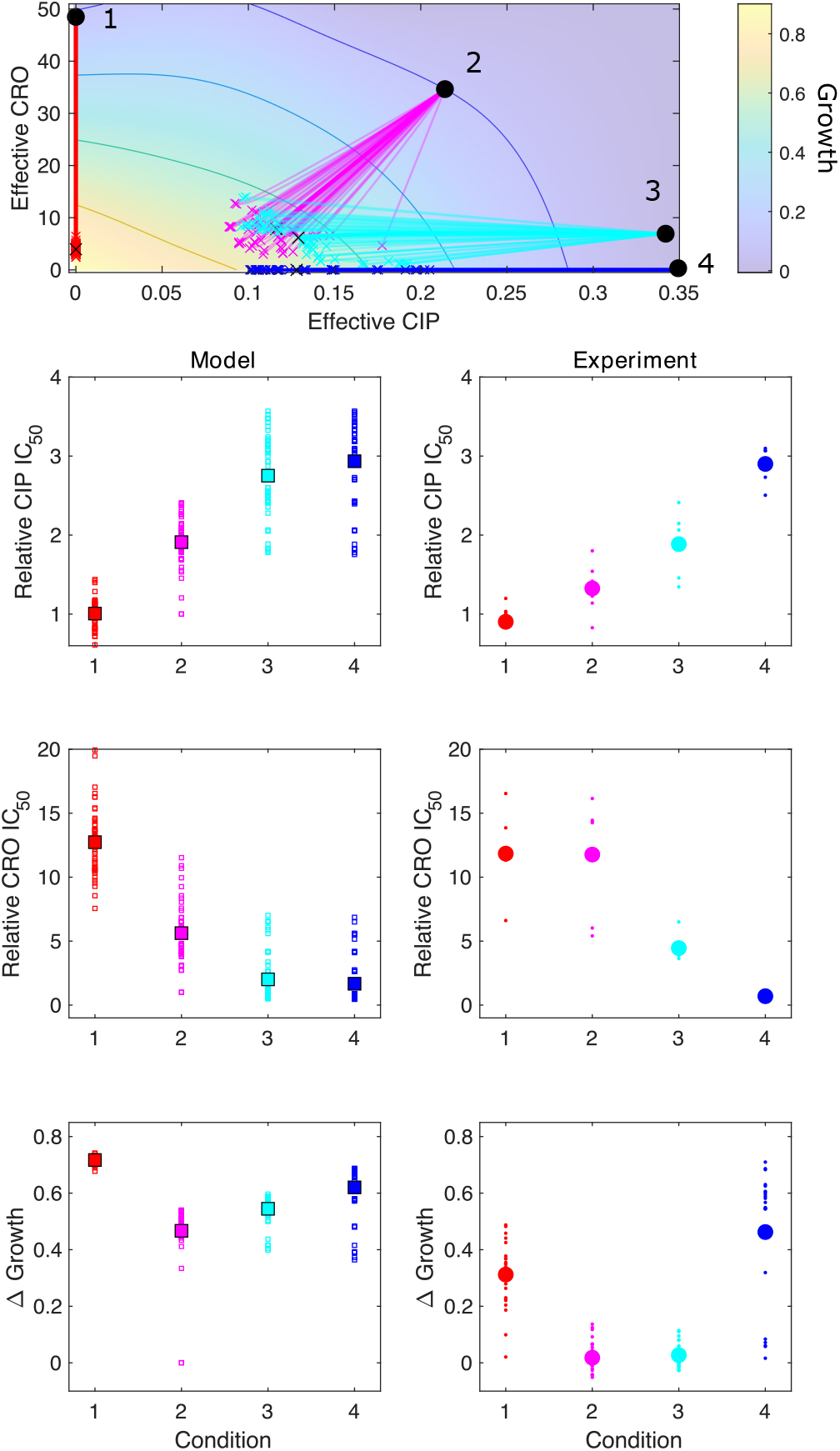
Model captures qualitative features of experimental evolution in combinations of CRO and CIP. Top panel: experimentally measured growth response surface for ceftriaxone (CRO) and ciprofloxacin (CIP). Circles represent 4 different adaptation conditions, each corresponding to a specific dosage combination (*x*_0_, *y*_0_). Solid lines show adaptation trajectories (i.e. changes in effective drug concentration over time) predicted from the Price Equation over an interval t=48 hrs. In each case, the set of available mutants–and hence, the set of possible scaling parameters *α* and *β*–is determined by sampling from experimentally measured changes in resistance (half-maximal inhibitory concentration IC_50_) for each drug in the full ensemble of mutants arising from adaptation to these drugs (see Figure S2). Bottom panels: comparing results from the model (left column) and experiment (right panel). Quantities include relative change in IC_50_ for each drug (top two rows) as well as overall change in population growth (bottom row);in each case, results are shown for each of the 4 conditions in the top panel. For the experiments, small markers are results from individual populations; larger markers are averages over all populations adapted to a given condition. For the model, small markers are results from each sub-sampling of the experimental scaling parameters;larger markers are averages over all sub-samples. Experimental data from (***Dean et al., 2020***).

**Figure S5.**
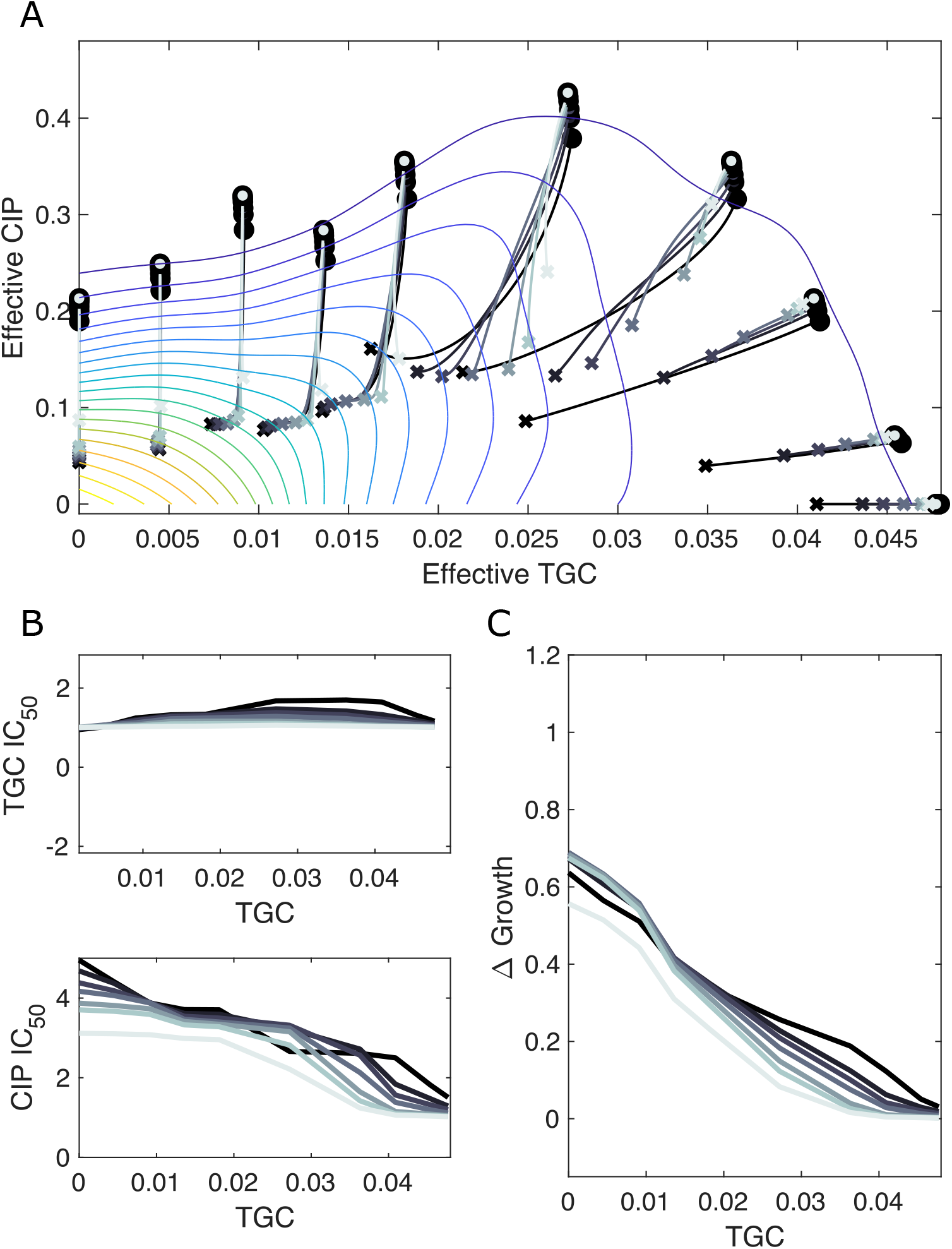
Resampling ensemble of scaling parameters does not dramatically impact dynamics. **A.** Experimentally measured growth response surface for tigecycline (TGC) and ciprofloxacin (CIP) from (***Dean et al., 2020***). Circles represent 11 different adaptation conditions, each corresponding to a specific dosage combination (*x*_0_, *y*_0_). Solid lines show the mean adaptation trajectories (i.e. changes in effective drug concentration over time) predicted from the Price Equation. In each case, the set of available mutants–and hence, the set of possible scaling parameters *α* and *β*-is determined by sampling *M* pairs from experimentally measured scaling parameters (from a total of 72 pairs). Different curves are averages over 100 realizations (samples) of *M* = 72 (darkest), *M* = 36, *M* = 24, *M* = 12, *M* = 7, *M* = 4, and *M* = 2 (lightest) *α_i_ − β_i_* pairs. In each realization, the ancestor frequency (*α = β* = 1) was fixed at 99% and the remaining 1% was randomly spread among the available mutants. **B.** Predicted change in IC_50_ for populations adapted in each of the 11 conditions in A. **C.** Predicted change in population growth rate for populations adapted in each condition in A.

**Figure S6.**
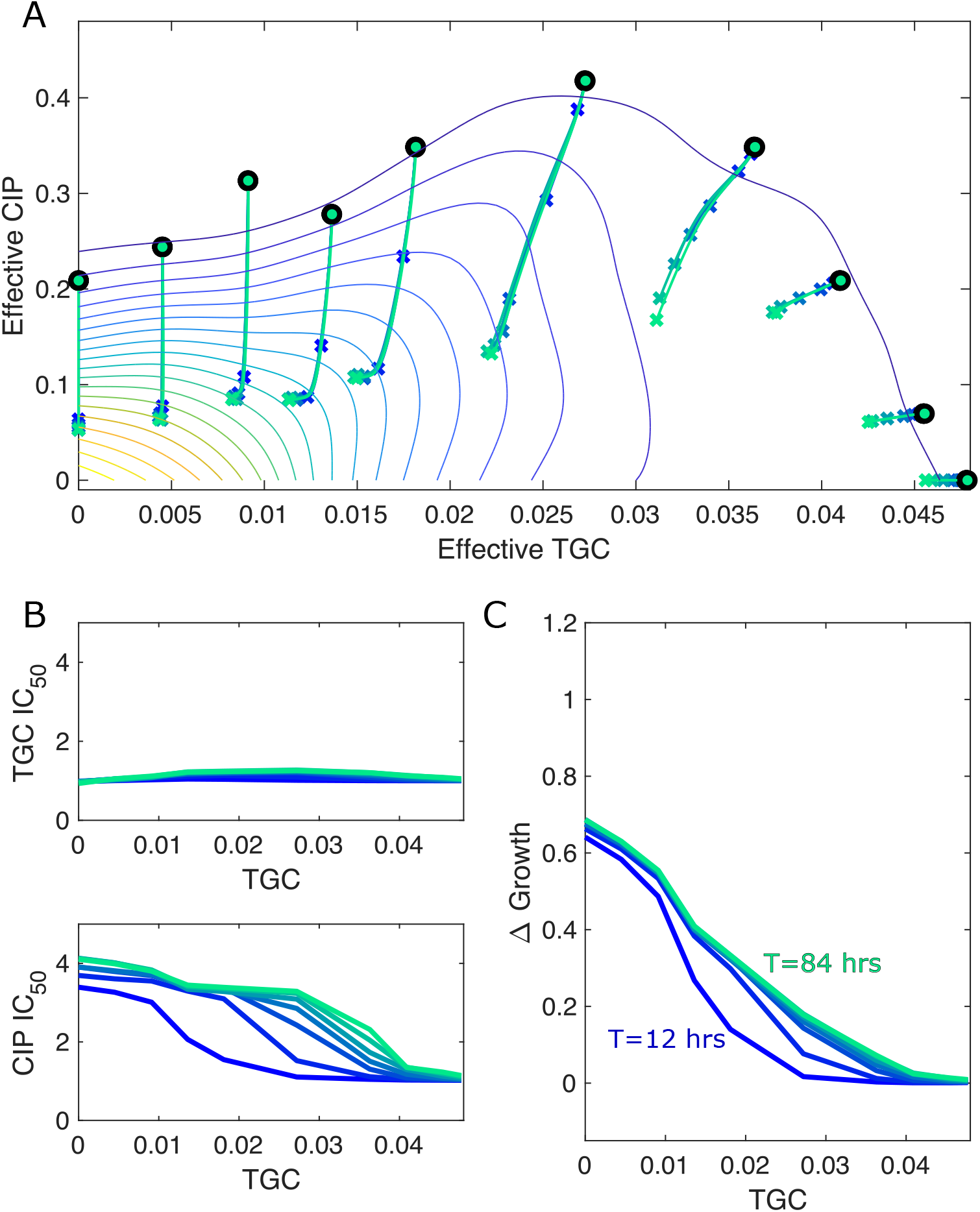
Different estimates of total time lead to similar qualitative dynamics. **A.** Experimentally measured growth response surface for tigecycline (TGC) and ciprofloxacin (CIP) from (***Dean et al., 2020***). Circles represent 11 different adaptation conditions, each corresponding to a specific dosage combination (*x*_0_, *y*_0_). Solid lines show the mean adaptation trajectories (i.e. changes in effective drug concentration over time) predicted from the Price Equation. In each case, the set of available mutants–and hence, the set of possible scaling parameters *α* and *β*–is determined by sampling *M* = 12 pairs from experimentally measured scaling parameters (from a total of 72 pairs). Different curves are averages over 100 realizations (samples) for total simulation times ranging from *T* = 12 to *T* = 84 hrs. In each realization, the ancestor frequency (*α* = *β* = 1) was fixed at 99% and the remaining 1% was randomly spread among the available mutants. **B.** Predicted change in IC_50_ for populations adapted in each of the 11 conditions in A. **C.** Predicted change in population growth rate for populations adapted in each condition in A. In panels B and C, the horizontal axis (TGC) is the external concentration of tigecycline (*μ*g/mL) used in each selecting condition in A.

**Figure S7.**
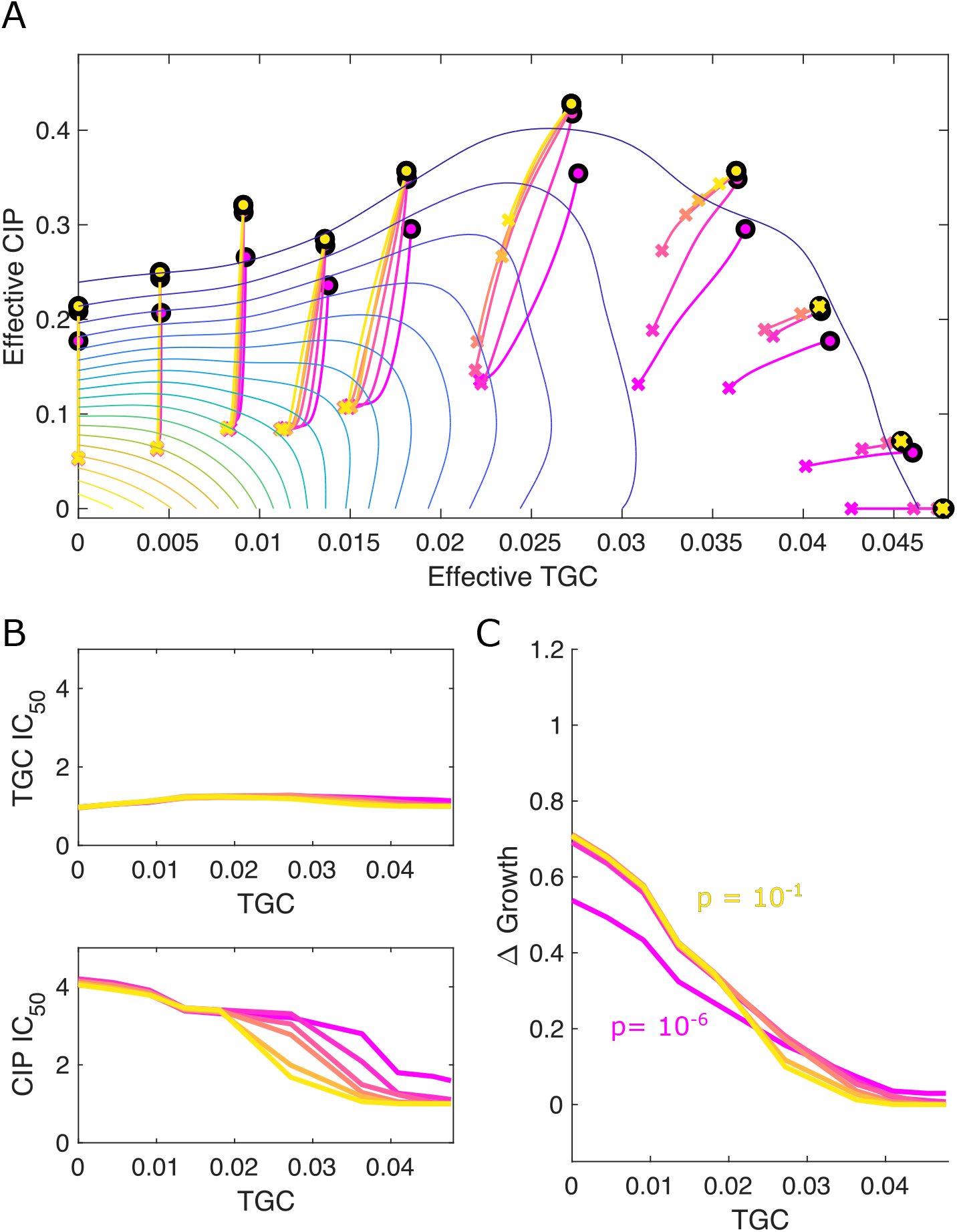
Different estimates of initial resistant fraction lead to similar qualitative dynamics. **A.** Experimentally measured growth response surface for tigecycline (TGC) and ciprofloxacin (CIP) from (***Dean et al., 2020***). Circles represent 11 different adaptation conditions, each corresponding to a specific dosage combination (*x*_0_, *y*_0_). Solid lines show the mean adaptation trajectories (i.e. changes in effective drug concentration over time) predicted from the Price Equation. In each case, the set of available mutants–and hence, the set of possible scaling parameters *α* and *β*–is determined by sampling *M* = 12 pairs from experimentally measured scaling parameters (from a total of 72 pairs). Different curves are averages over 100 realizations (samples) for initial mutant fractions ranging from *p* = 10^−1^ to *p* = 10^−6^ hrs. In each realization, the dynamics were simulated for a total time of *T* = 72 hrs **B.** Predicted change in IC_50_ for populations adapted in each of the 11 conditions in A. **C.** Predicted change in population growth rate for populations adapted in each condition in A. In panels B and C, the horizontal axis (TGC) is the external concentration of tigecycline (*μ*g/mL) used in each selecting condition in A.

